# Horizontal gene transfer in the human and skin commensal *Malassezia*: a bacterially-derived flavohemoglobin is required for NO resistance and host interaction

**DOI:** 10.1101/2020.01.28.923367

**Authors:** Giuseppe Ianiri, Marco A. Coelho, Fiorella Ruchti, Florian Sparber, Timothy J. McMahon, Ci Fu, Madison Bolejack, Olivia Donovan, Hayden Smutney, Peter Myler, Fred Dietrich, David Fox, Salomé LeibundGut-Landmann, Joseph Heitman

## Abstract

The skin of humans and animals is colonized by commensal and pathogenic fungi and bacteria that share this ecological niche and have established microbial interactions. *Malassezia* are the most abundant fungal skin inhabitant of warm-blooded animals, and have been implicated in skin diseases and systemic disorders, including Crohn’s disease and pancreatic cancer. Flavohemoglobin is a key enzyme involved in microbial nitrosative stress resistance and nitric oxide degradation. Comparative genomics and phylogenetic analyses within the *Malassezia* genus revealed that flavohemoglobin-encoding genes were acquired through independent horizontal gene transfer events from different donor bacteria that are part of the mammalian microbiome. Through targeted gene deletion and functional complementation in *M. sympodialis*, we demonstrated that bacterially-derived flavohemoglobins are cytoplasmic proteins required for nitric oxide detoxification and nitrosative stress resistance under aerobic conditions. RNAseq analysis revealed that endogenous accumulation of nitric oxide resulted in upregulation of genes involved in stress response, and downregulation of the MalaS7 allergen-encoding genes. Solution of the high-resolution X-ray crystal structure of *Malassezia* flavohemoglobin revealed features conserved with both bacterial and fungal flavohemoglobins. *In vivo* pathogenesis is independent of *Malassezia* flavohemoglobin. Lastly, we identified additional 30 genus- and species-specific horizontal gene transfer candidates that might have contributed to the evolution of this genus as the most common inhabitants of animal skin.

**Significance statement:** *Malassezia* species are the main fungal components of the mammalian skin microbiome and are associated with a number of skin disorders. Recently, *Malassezia* has also been found in association with Crohn’s Disease and with pancreatic cancer. The elucidation of the molecular bases of skin adaptation by *Malassezia* is critical to understand its role as commensal and pathogen. In this study we employed evolutionary, molecular, biochemical, and structural analyses to demonstrate that the bacterially-derived flavohemoglobins acquired by *Malassezia* through horizontal gene transfer resulted in a gain of function critical for nitric oxide detoxification and resistance to nitrosative stress. Our study underscores horizontal gene transfer as an important force modulating *Malassezia* evolution and niche adaptation.

## Introduction

The skin microbiome includes numerous microorganisms that establish a variety of direct and indirect interactions characterized by the exchange of genetic material that impact microbial biology contributing to their speciation and evolution. *Malassezia* is the most abundant fungal genus resident on human skin, representing more than 90% of the skin mycobiome (1). This genus presently consists of 18 diverse species (2), each with an unusually compact genome that underwent extensive gene turnover events as a result of evolutionary adaptation and colonization to a nutrient-limited ecological niche such as the skin (3). Although commensals, *Malassezia* species are also associated with a number of clinical skin disorders, including pityriasis versicolor, dandruff, and atopic dermatitis (AD) (4). A recently-developed epicutaneous murine model revealed that the host responses to *Malassezia* are dominated by pro-inflammatory cytokine IL-17 and related factors that prevent fungal overgrowth and exacerbate inflammation under atopy-like conditions (5). Furthermore, *Malassezia* species have also been implicated recently as causal agents of Crohn’s Disease/Inflammatory Bowel Disease in patients with *CARD9* mutations, and in accelerating the progression of Pancreatic Adenocarcinoma in murine models and in humans, and cystic fibrosis pulmonary exacerbation (6–8).

Nitric Oxide (NO) is a reactive compound of central importance in biological systems and it functions both as a signaling and toxic molecule. While little is known about NO synthesis in fungi, in mammals NO is synthesized by NO synthases (NOS isoforms). Nos1 and Nos3 are constitutively expressed in neurons and endothelium, respectively, and produce NO to promote S-nitrosylation and transcriptional regulation. S-nitrosylation is a post-translational mechanism involving oxidative modification of cysteine by NO, and this is the central NO-mediated signaling mechanism that affects myriad of cellular physiological and pathophysiological processes (9). On the other hand, the Nos2 is not constitutively expressed but is induced in inflammatory cells in response to infection and is involved in wound healing, immune regulation, and host defense (10).

In fungi, NO is synthesized through a reductive denitrification pathway from nitrite, and through an oxidative pathway from L-arginine, although the detailed biochemical mechanisms have not yet been fully elucidated (11–13). Compared to mammals, plants, and bacteria, the role of NO in fungal biology is understudied. In *S. cerevisiae* NO is important for activation of transcription factors that are involved in resistance to a variety of environmental stress conditions, such as oxidative stress, heat shock, and hydrostatic pressure (11). Other studies report an involvement of NO in pathogenesis of *Botrytis cinerea* and *Magnaporthe oryzae*, in morphogenesis and reproduction in *Aspergillus nidulans*, and in the yeast-to-hyphae dimorphic transition in *Candida albicans* (12, 14, 15).

Imbalance in cellular NO levels leads to altered redox homeostasis, resulting in the production of reactive nitrogen species that are responsible for nitrosative stress (10). NO dioxygenases are enzymes that living cells use to actively consume poisonous NO by converting it to inert nitrate, a source of nitrogen (16). Red blood cell hemoglobin is the main mammalian dioxygenase that metabolizes NO in the vascular lumen, whereas a type I flavohemoglobin constitutes the main enzyme deployed by microbes to counteract NO toxicity (10, 11). Some fungi within the *Aspergillus* genus have two type I flavohemoglobins, one that is cytosolic and protects the cells against exogenous NO, and another that is mitochondrial and is putatively involved in detoxification of NO derived from nitrite metabolism (17, 18). A type II flavohemoglobin has also been identified in *Mycobacterium tuberculosis* and other actinobacteria, but it lacks NO consuming activity and it utilizes D-lactate as an electron donor to mediate electron transfer (19, 20).

Evolution of flavohemoglobins in microbes has been previously investigated, revealing a dynamic distribution across bacteria and eukaryotes characterized by frequent gene loss, gene duplication, and horizontal gene transfer (HGT) events (21–23). An interesting finding of these phylogenetic studies was the HGT-mediated acquisition of a bacterial flavohemoglobin-encoding gene, *YHB1*, by *M. globosa* and *M. sympodialis* (21, 22). Here, we employed evolutionary, molecular, biochemical, and structural analyses to demonstrate that the HGT of the bacterial flavohemoglobin in *Malassezia* resulted in a gain of function critical for resistance to nitrosative stress and detoxification of NO under aerobic conditions. Moreover, analysis of the available *Malassezia* genomes revealed that extant flavohemoglobin-encoding genes are present as a single copy in different species, and resulted from a complex pattern of differential retention/loss of one of two genes (*YHB1* and *YHB101*), which were ancestrally acquired by *Malassezia* through HGT from different donor bacterial lineages. Through trans-species complementation we demonstrated that the second bacterially-derived flavohemoglobin identified, Yhb101, restores resistance of the *yhb1Δ* mutant to nitrosative stress-inducing agents and also is able to consume NO. RNAseq analysis revealed that endogenous accumulation of NO results in upregulation of genes involved in stress responses and transport, and downregulation of the allergen-encoding genes. The characterization of the X-ray crystal structure of the *Malassezia* flavohemoglobin revealed features shared with bacterial and fungal flavohemoglobins. Moreover, *ex vivo* and *in vivo* experiments suggest that *Malassezia* flavohemoglobins are dispensable for pathogenesis under our tested experimental conditions. Because the HGT-mediated acquisition of the flavohemoglobin-encoding genes conferred the ability to metabolize NO, *Malassezia* genomes were searched for other HGT that could represent important gain of function events. Thirty additional genus- and species-specific HGT events were identified, with the donors being predominantly Actinobacteria and Proteobacteria. Similar to *Malassezia*, these donor species are some of the most common members of human and mammalian microbiomes, suggesting that niche overlap may have enhanced the opportunity for inter-kingdom HGT.

## Results

### *Malassezia* flavohemoglobin-encoding genes were ancestrally acquired from bacteria through independent HGT events

Flavohemoglobins are critical for nitric oxide (NO) detoxification and counteract nitrosative stress (10). Previous studies reported that the flavohemoglobin-encoding gene *YHB1* was acquired by *M. globosa* and *M. sympodialis* through HGT from *Corynebacterium*, a bacterial genus within the Actinobacteria that includes species that are part of the human microbiome (21, 22).

Because flavohemoglobins are widespread in both bacteria and eukaryotes, we examined whether the remaining 13 sequenced *Malassezia* species also contain a flavohemoglobin-encoding gene, and whether it had a fungal or bacterial origin. BLAST analyses with the *M. globosa* Yhb1 sequence as query identified a single copy of *YHB1* in all *Malassezia* species and strains with sequenced genomes. Intriguingly, this comparative search revealed that the best hits in *M. yamatoensis* and *M. slooffiae* had lower *E*-values (5e-64 and 1e-59, respectively) compared to the remaining *Malassezia* species (*E*-values ranging from 0.0 to 7e-161), possibly suggesting different origins or modifications of these flavohemoglobin-encoding genes. To elucidate the evolutionary trajectory of flavohemoglobin within the whole *Malassezia* genus, a Maximum Likelihood (ML) phylogenetic tree was reconstructed using >2000 Yhb1 bacterial and fungal sequences retrieved from GenBank. This phylogenetic analysis revealed two clades of flavohemoglobin within *Malassezia* genus: clade 1 that includes 13 species and clusters together with *Brevibacterium* species belonging to Actinobacteria; and clade 2 that includes *M. yamatoensis* and *M. slooffiae* and clusters together with different Actinobacteria with the closest relative being *Kocuria kristinae* (Fig. 1 A-C). To evaluate the statistical phylogenetic support for the two *Malassezia* flavohemoglobin clades whose distribution was not monophyletic, we performed approximately unbiased (AU) comparative topology tests. The constrained ML phylogeny, in which all *Malassezia* flavohemoglobins were forced to be monophyletic was significantly rejected (AU test, *P* value = 0.001, Fig. 1D), thus not supporting the null hypothesis that all flavohemoglobin genes in *Malassezia* have a single origin.

**Figure 1.**
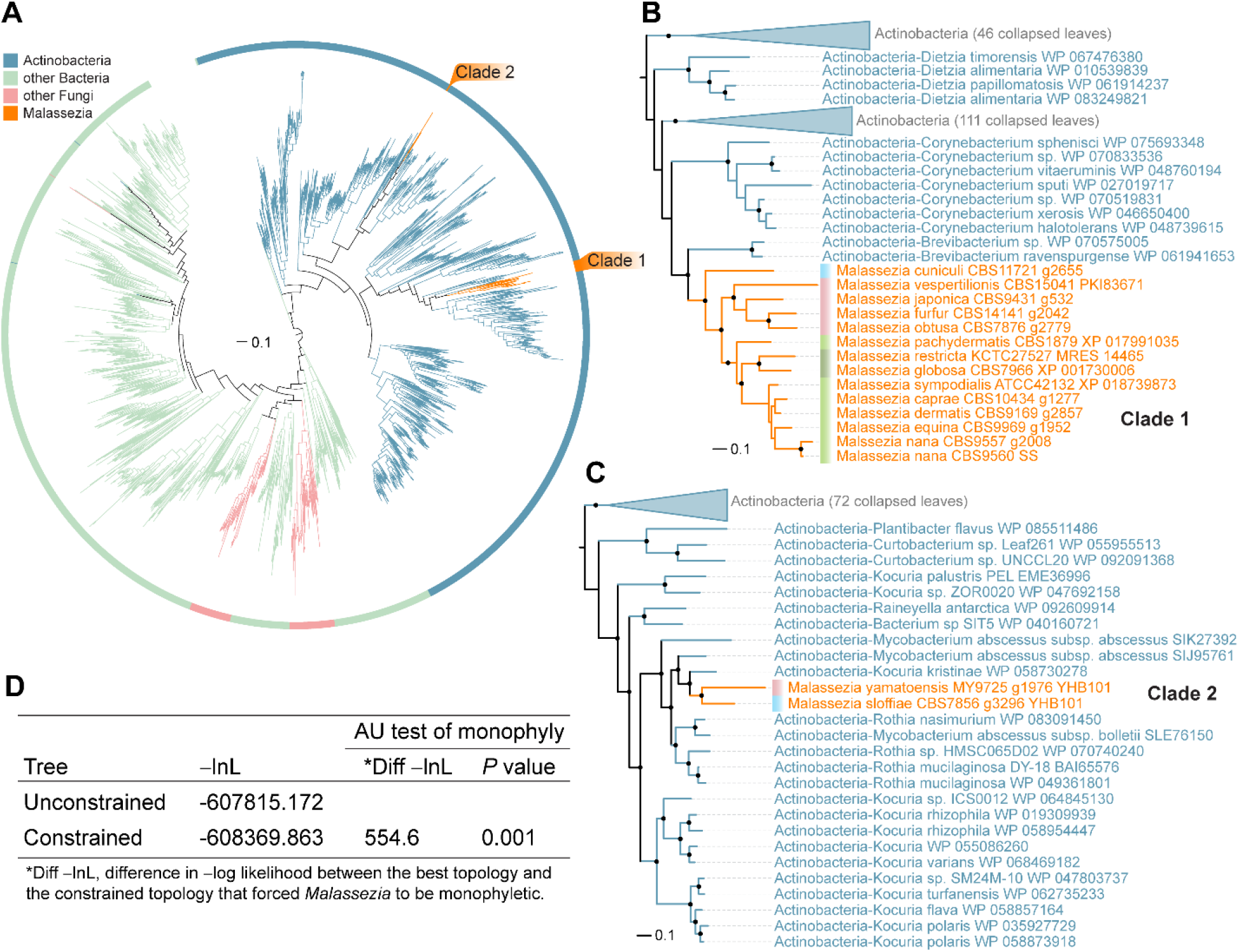
Evidence for independent HGT events of the flavohemoglobin-encoding genes in *Malassezia* from the Actinobacteria. (A) Maximum likelihood phylogeny of consisting of 2,155 flavohemoglobin protein sequences. Two groups (clade 1 and 2) of horizontally transferred flavohemoglobin genes (*YHB1* and *YHB101*) in *Malassezia* are colored in orange. Other tree branches are colored according to the key on the top left, representing other major groups of organisms. The phylogeny was visualized using iTOL v3.6.1 (98) and was rooted at the midpoint. (B and C) Zoomed views of the ML phylogeny showing in more detail the position of *Malassezia* flavohemoglobins from clades 1 and 2, and their putative bacterial donor lineages. (D) Results of topological constraints tests that significantly rejected the monophyletic origin for both *Malassezia* flavohemoglobins clades, providing additional support for independent HGT events.

To validate this further, a region of ~30 kb surrounding the flavohemoglobin encoding gene in all sequenced *Malassezia* species was subjected to synteny comparison (Fig. 2). Overall, this region was highly syntenic across species, with the exception of some lineage-specific rearrangements located upstream of *YHB1* (Fig. 2B). Remarkably, while the same regions were also highly conserved in *M. yamatoensis* and *M. slooffiae*, they both lacked the flavohemoglobin-encoding gene, which is instead located in a different, non-syntenic, regions of their genomes (Fig. 2B and Fig. S1). Therefore, both phylogenic and synteny comparisons strongly support the hypothesis that *Malassezia* flavohemoglobin genes were acquired through independent HGT events from different bacterial donor species. We named the flavohemoglobin of clade 1 Yhb1 following the *S. cerevisiae* nomenclature (24), and that of *M. yamatoensis* and *M. slooffiae* Yhb101 (clade 2). The two different flavohemoglobins protein sequences share 38% identity (Fig. S2).

**Figure 2.**
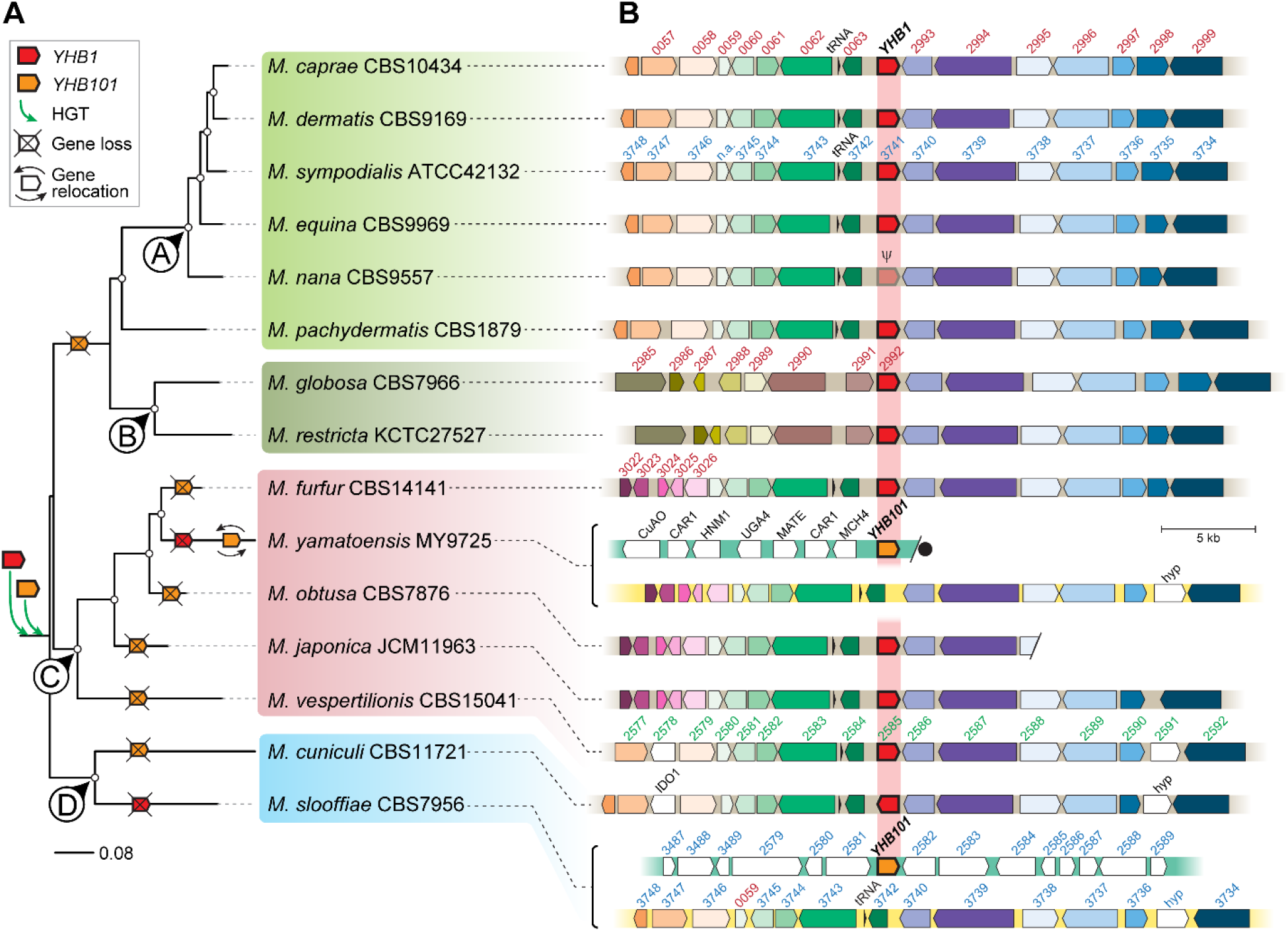
Evolutionary trajectory of flavohemoglobin-encoding genes in *Malassezia* after their acquisition via HGT from different donor bacteria lineages. (A) Phylogenetic relationship of *Malassezia* species with available genome sequence inferred from the concatenation of 246 single-copy proteins. Color codes assigned to the different phylogenetic clades (named A to D) are kept consistent in all figures. The tree was rooted at the midpoint and white circles in the tree nodes indicate full UFboot and SH-aLRT branch support. The proposed evolutionary events that led to the final arrangement of the flavohemoglobin-encoding genes shown in panel B are shown in the phylogenetic tree, as given in the key; double arrows indicate relocation of the *YHB101* gene in subtelomeric position. (B) Chromosomal regions encompassing the *YHB1* gene in *Malassezia*. Genes are shown as arrows denoting the direction of transcription and orthologs are represented in the same color. Non-syntenic genes are shown in white, and small arrows in black represent tRNAs. The *YHB1* gene is shown as red arrows outlined in bold in the center. The end of a scaffold is represented by a forward slash. For *M. yamatoensis* and *M. slooffiae*, yellow bars indicate the absence of the *YHB1* gene in otherwise syntenic regions, and those in green indicate instances where another flavohemoglobin-encoding gene, named *YHB101* and represented as orange arrows outlined in bold, was acquired by an independent HGT event. A defective *YHB1* gene in *M. nana* CBS9557 is denoted by the Greek symbol ψ. Gene codes in red or blue are as they appear in *M. globosa* (prefix “MGL_”) or *M. sympodialis* (prefix “MSYG_”) genome annotations, respectively, those in black were named based on top BLASTp hits in *S. cerevisiae*, and “hyp” represent hypothetical proteins. Black circle represents the end of a chromosome. Scaffold/chromosomal locations and accession numbers are given for each region in Table S1.

Because several studies suggest that genomic regions flanking horizontally-acquired genes are enriched in DNA transposons and retrotransposons (25, 26), a 5 kb region surrounding the flavohemoglobin-encoding genes was analyzed in two *Malassezia* species representative of clade1 (*M. sympodialis*) and clade 2 (*M. slooffiae*). Dot plot comparisons revealed overall high co-linearity with the common flanking genes encoding a hypothetical protein and Nsr1 (Fig. S3A). Interestingly, a highly repetitive sequence that shares similarity with the long terminal repeat (LTR) Gypsy was identified in the *NSR1* gene flanking *YHB1* (Fig. S3A-B), and we speculate that this LTR-like region might have facilitated the non-homologous end joining (NHEJ) integration of the bacterial *YHB1* gene into the *Malassezia* common ancestor.

The flavohemoglobin-encoding gene *YHB101* in *M. yamatoensis* and *M. slooffiae* seems to have been acquired in a single HGT event from the same bacterial donor lineage (Fig. 1A-C), although its genomic location is not syntenic in these two species (Fig. 2B). In *M. slooffiae*, *YHB101* is located in a region that is otherwise highly conserved across *Malassezia* (Fig. S1A-B) and is devoid of transposable elements or repetitive regions that could have facilitated NHEJ of *YHB101* (Fig. S1B). In contrast, in *M. yamatoensis YHB101* is located at the end of a chromosome and the adjacent genes are not syntenic in other *Malassezia* species, with the exception of a more distant group of five genes (from *JLP1* to *MSS1*) (Fig. S1C). In other *Malassezia* species (e.g. *M. japonica, M. slooffiae, M. sympodialis*) these five genes are subtelomeric, suggesting that chromosomal reshuffling might have contributed to generate the unique arrangement of genes surrounding *M. yamatoensis YHB101* (Fig. 2; Fig. S1C).

Based on the analyses performed and on the availability of *Malassezia* genomes, we propose the following evolutionary model of flavohemoglobin-mediated HGT in *Malassezia*. First, the *YHB1* and *YHB101* genes were independently acquired by the *Malassezia* common ancestor via HGT from a *Brevibacterium*-related and a *Kocuria*-related bacterial donor, respectively. An early loss of the *YHB101* subsequently occurred in the common ancestor of the lineages that include *M. sympodialis* and *M. globosa* (Clades A and B, Fig. 2A), which retained the *YHB1* gene in its ancestral location; lack of synteny in the region upstream of the *YHB1* gene in *M. globosa* and *M. restricta* represents a more recent chromosomal rearrangement (Fig. 2B). In the early-branching *Malassezia* lineage (Clade D, Fig. 2A), the *YHB1* gene was lost in *M. slooffiae*, which instead retained the *YHB101* gene, presumably in its ancestral location; conversely, *M. cuniculi* lost the *YHB101* gene and retained the *YHB1* gene in the ancestral location. Lastly, all species within the *M. furfur* lineage (Clade C, Fig. 2A) have the *YHB1* gene in its ancestral location, with the exception of *M. yamatoensis* that has lost this gene and retained instead the *YHB101* gene, which was then relocated from its original position to a subtelomeric region (Fig. 2B). This model implies that *M. vespertilionis, M. japonica, M. obtusa*, and *M. furfur* have independently lost the *YHB101* gene during their evolution (Fig. 2A). Because none of the *Malassezia* species has the two flavohemoglobin genes (*YHB1* and *YHB101*) in their genomes, we posit that loss of one or the other flavohemoglobin may be a consequence of different selection pressures across descendant lineages of the HGT recipient.

### Bacterially-derived flavohemoglobin-encoding genes are required for nitrosative stress resistance and NO detoxification in *Malassezia*

Flavohemoglobins are critical for NO detoxification and counteract nitrosative stress (10). To assess whether this HGT event in *Malassezia* resulted in a gain of function, we deleted the *YHB1* ORF (*MSYG_3741*) of *M. sympodialis* ATCC42132 through targeted mutagenesis using our recently developed transformation protocol based on transconjugation mediated by *Agrobacterium tumefaciens* (27, 28) (Fig. S4A).

The *M. sympodialis yhb1*Δ mutant exhibits hypersensitivity to the NO-donors DETA NONOate and sodium nitrite (NaNO_2_), but not to hydrogen peroxide (H_2_O_2_) (Fig. 3A). The two identified *Malassezia* flavohemoglobins Yhb1 and Yhb101 were used to generate GFP fusion proteins whose expression was driven by the respective endogenous promoter to complement the *M. sympodialis yhb1*Δ mutant phenotype and to assess protein localization (Fig. S4B). Reintroduction of either flavohemoglobin in the *M. sympodialis yhb1*Δ mutant restored resistance to nitrosative stress at the WT level (Fig. 3A). In agreement, fusion protein expression in complemented strains was confirmed by qPCR (Fig. S4C-D), FACS, and fluorescence microcopy imaging of GFP expression, which revealed that *Malassezia* flavohemoglobins are cytoplasmic (Fig. 3B).

**Figure 3.**
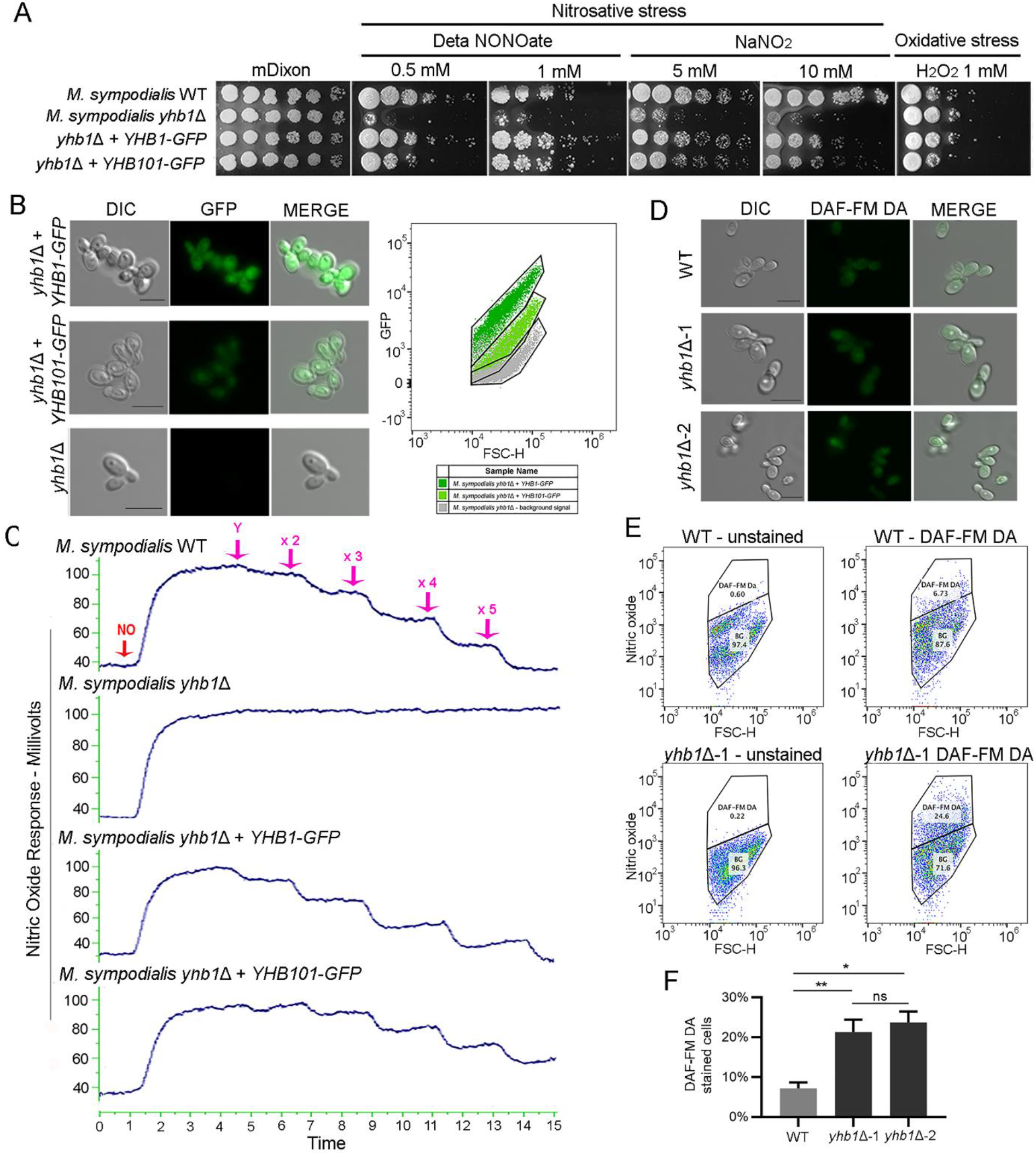
*M. sympodialis* flavohemoglobins are involved in nitrosative stress resistance and NO degradation. (A) Stress sensitivity assay of *M. sympodialis* WT, *yhb1*Δ mutant, and complementing strains *yhb1*Δ + *YHB1* and *yhb1*Δ + *YHB101* on mDixon agar supplemented with the NO-donor agent DETA NONOate and NaNO2, and with hydrogen peroxide. (B) GFP expression in the *M. sympodialis yhb1*Δ mutant, and complementing strains *yhb1*Δ + *YHB1* and *yhb1Δ* + *YHB101*, and respective GFP signal analyzed trough FACS. (C) NO consumption assay by *M. sympodialis* WT, *yhb1Δ* mutant, and complementing strains *yhb1Δ* + *YHB1* and *yhb1Δ* + *YHB101;* the blue trace indicates the NO level over a period of 15 min. NO and *Malassezia* (Y = yeast) injections are indicated by purple arrows. In this experiment, 10 μL (Y), 20 μL (x 2) and 30 μL (x 3), 40 μL (x 4) and 50 μL (x 5) of *Malassezia* cellular suspensions were injected. (D) Representative fluorescent staining of intracellular NO with DAF-FM DA in *M. sympodialis* WT and two independent *yhb1Δ* mutants, and (E-F) quantification of the NO signal by flow cytometry; spontaneous fluorescence of *M. sympodialis* was used as background to detect specifically DAF-FM DA signal. Asterisks indicates statistically significant differences (* = p<0.05, ** = p<0.01) according to the unpaired student’s t-test with Welch’s correction.

Next, we tested whether *Malassezia* flavohemoglobins were able to actively detoxify NO, which could potentially account for its involvement in nitrosative stress resistance. To this aim, we adapted a biochemical assay used for evaluating NO consumption by hemoglobin in red blood cells and plasma (29) to the commensal yeast *Malassezia* (Fig. S5). As shown in Figure 2C, while the *M. sympodialis* WT strain exhibited robust and dose-dependent NO degradation, the *yhb1Δ* mutant showed no NO consumption. Complemented strains were able to actively consume NO, although *M. sympodialis yhb1Δ* + *YHB101-GFP* displayed a lower NO consumption, corroborating the results of the phenotypic assay, GFP expression, and FACS analysis (Fig. 3A-C). These differences might be due to less efficient cross-species complementation of the *M. yamatoensis* Yhb101-GFP fusion protein in the *yhb1*Δ mutant of *M. sympodialis*, which was a strategy chosen because of the lack of protocols for gene deletion in *M. slooffiae* and *M. yamatoensis*. Taken together these genetic and biochemical analyses show that the bacterially-derived flavohemoglobins protect *Malassezia* from nitrosative stress by decreasing toxic levels of NO.

To assess intracellular production of NO by *M. sympodialis*, cells were stained with the NO-specific dye 4-Amino-5-methylamino-2’,7’-diaminofluorescein diacetate (DAF-FM DA), which passively diffuses across membranes and emits increased fluorescence after reacting with NO. Fluorescent microscopy revealed intracellular accumulation of NO in both the WT and *yhb1Δ* mutant of *M. sympodialis* (Fig. 3D). NO-staining was quantified by FACS analysis, revealing significantly higher NO accumulation in the flavohemoglobin mutant *yhb1Δ* compared to the *M. sympodialis* WT (Fig. 3E-F). Because DAF-FM DA and GFP have similar excitation/emission spectra, complemented strains could not be tested for NO accumulation via flow cytometry, and therefore an independent *M. sympodialis yhb1Δ* mutant was tested and yielded similar results. These results indicate that the lack of a functional flavohemoglobin leads to intracellular accumulation of NO.

Finally, a broader analysis was performed to assess other functions of the *Malassezia* flavohemoglobins in response to a variety of environmental stresses and clinical antifungals, but in all cases the *M. sympodialis yhb1*Δ mutant phenotype was not significantly different from the WT (Fig. S6). These phenotypic results are in agreement with those obtained for the basidiomycetous yeast *Cryptococcus neoformans* (30, 31), but contrast with the studies carried out in ascomycetous fungi, in which in addition to NO and nitrosative stress sensitivity, flavohemoglobin mutants exhibited higher resistance to hydrogen peroxide in *A. nidulans* (32), and hyper-filamentation in *C. albicans* (33). Several studies also report the protective role of both bacterial and fungal flavohemoglobins against NO under anaerobic conditions (10, 24, 34). However, we could not confirm this function for the *Malassezia* flavohemoglobin in our anaerobic experiments because no phenotypic differences were observed between the WT, the*yhb1Δ* mutant, and the complemented strains (Fig. S7).

### A recent inactivation of *YHB1* in *M. nana* results in compromised NO enzymatic consumption

Analysis of Yhb1 protein prediction across species revealed that *M. nana YHB1* underwent pseudogenization [i.e. loss of gene function by disruption of its coding sequence with generation of a pseudogene, which is usually indicated as ψ (35)] following a G-to-T transversion in the glycine codon GGA, generating a premature TGA stop codon at the 29^th^ amino acid (Fig. S8A). Literature search revealed that the sequenced strain of *M. nana* CBS9557 was isolated in Japan from a cat with otitis externa (36), while the other four known *M. nana* strains were collected in Brazil: *M. nana* CBS9558 and CBS9559 from cows with otitis externa, and CBS9560 and CBS9561 from healthy cows (36).

The *M. nana* strains CBS9557, CBS9559, and CBS9560 were used to investigate whether the pseudogenization event occurred in a *M. nana* ancestor, and whether it impacts nitrosative stress resistance and NO consumption. Because no genomes are available for the *M. nana* strains CBS9559 and CBS9560, their *YHB1* gene was amplified by PCR and Sanger sequenced using primers designed on the *YHB1* of *M. nana* CBS9557 (Table S2). *YHB1* sequence comparison confirmed a premature stop codon present in only CBS9557, with both Brazilian *M. nana* isolates having a full-coding *YHB1* gene (Fig. S8A). Phenotypic analysis revealed no significant difference in resistance to nitrosative stress by the three *M. nana* strains, with only a modest increased sensitivity displayed by CBS9557 exposed to 10 mM of sodium nitrite (Fig. S8B). Strikingly, *M. nana* CBS9557 displayed undetectable NO consumption activity as observed for the *M. sympodialis yhb1Δ* mutant, while *M. nana* CBS9559 and CBS9560 showed regular dose-dependent NO consumption (Fig. S8C). These data suggest that the inactivation of flavohemoglobin in *M. nana* CBS9557 impaired the ability to consume NO, but this does not impact the resistance to nitrosative stress, which might be compensated by other stress responsive pathways.

Intriguingly, another pseudogenization event of a bacterial gene encoding an aliphatic amidase was also identified in *M. nana* CBS9557 (represented in Fig. 6). These nonsense mutations were identified only in the *M. nana* CBS9557 isolated in Japan, suggesting that the different origin of the *M. nana* strains might contribute to this intraspecies diversity. This hypothesis is further supported by different phenotypic traits displayed by the *M. nana* isolates (Fig. S9). Exposure to several stress conditions revealed different responses to the most common antifungal drugs by *M. nana* strains, with strain CBS9557 displaying increased sensitivity to amphotericin B and resistance to fluconazole, and the geographically-related strains CBS9559 and CBS9560 displaying an opposite phenotype (Fig. S9).

### NO accumulation in *M. sympodialis* leads to upregulation of genes involved in nitrogen metabolism, ergosterol biosynthesis, and protein folding, and downregulation of predicted pathogenicity factors

Because the *M. sympodialis* flavohemoglobin mutant *yhb1Δ* accumulates higher amounts of NO than the WT (Fig. 3 D-E), we compared their transcriptomic profile to elucidate any potential signaling role of endogenous NO. RNAseq analysis revealed 36 differentially expressed genes for false discovery rate (FDR) <0.05, of which 14 were upregulated and 22 were downregulated; using an additional threshold of log_2_FC +/-0.5 we found 3 upregulated and 9 downregulated genes (Fig. 4A; Dataset S1-S2). Of these, the only upregulated gene with log_2_FC > 1 encodes an uncharacterized protein (*MSYG_1280*), while two others with 0.5 < log_2_FC < 1 encode Nop56 (or Sik1), a nucleolar protein involved in pre-rRNA processing, and an uncharacterized protein (*MSYG_0148*) predicted to be involved in magnesium transport. Other known upregulated genes with log_2_FC <0.5 are involved in response to stresses and transport (Dataset S1). The majority of the downregulated genes include those encoding hypothetical proteins (5 out of 9), the regulator of phospholipase D Srf1, two MalaS7 allergens, and an uncharacterized allergen (Dataset S2). It is worth noting that a large number of differentially expressed genes (DEGs) are predicted to encode unknown proteins, suggesting novel and unknown signaling pathways regulated by endogenous NO in *Malassezia* (Dataset S1-S2).

**Figure 4.**
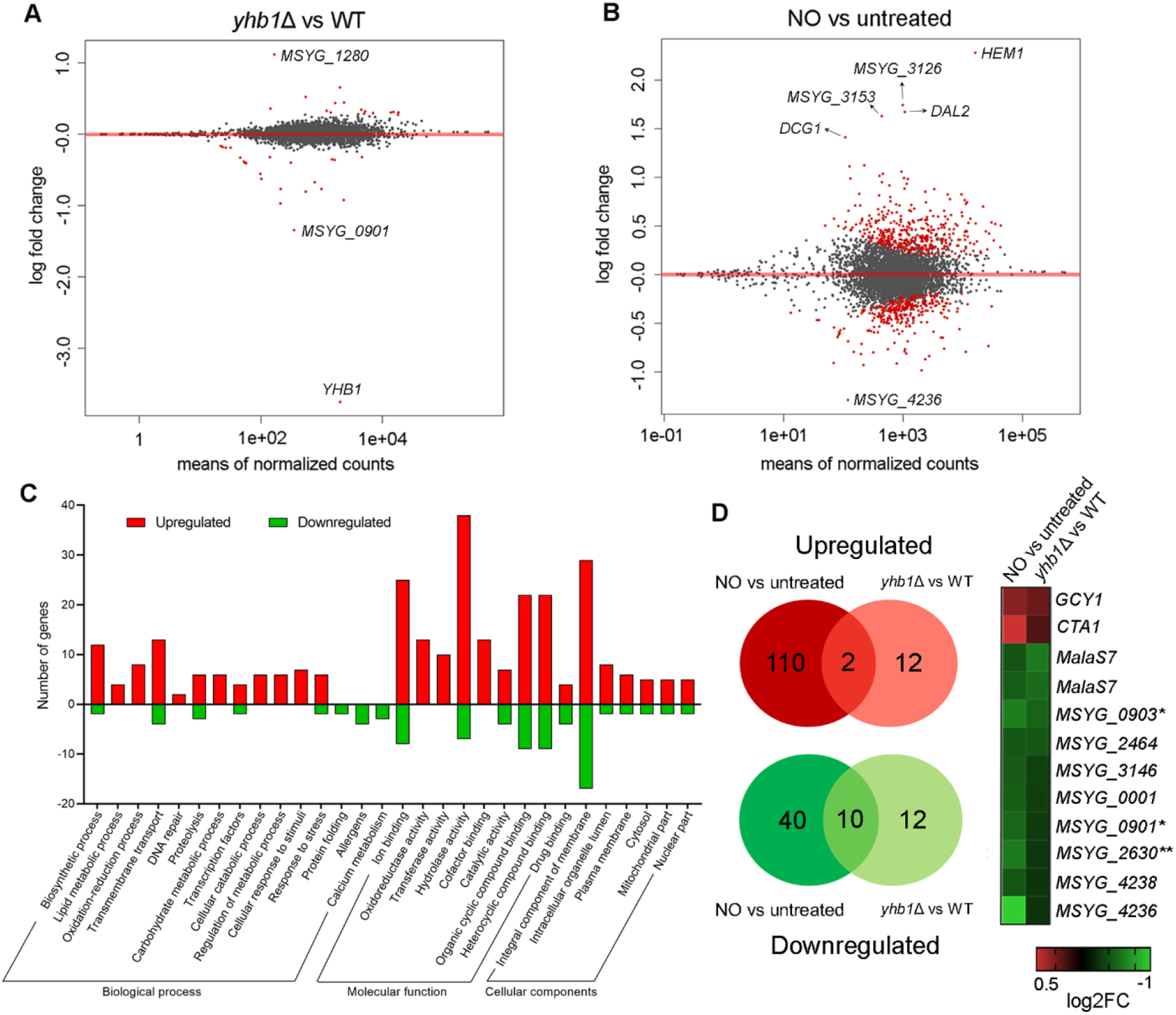
Transcriptomic profile of *M. sympodialis* strains under NO-accumulation conditions. (A) MA-plot displaying the transcriptomic changes of the *M. sympodialis yhb1*Δ mutant compared to the WT *M. sympodialis* strain. Red dots indicate differentially expressed genes for FDR < 0.05. The most upregulated and downregulated genes (*MSYG_1280* and*MSYG_0901*, respectively) are indicated, along with the *YHB1* gene, which represents an internal control as its downregulation is expected because the gene is deleted. (B) MA-plot displaying the transcriptomic changes of*M. sympodialis* WT grown in the presence of NaNO2 compared to the untreated control. Red dots indicate differentially expressed genes for FDR < 0.05; the most upregulated and downregulated genes are indicated. (C) Gene ontology classification relative to the RNAseq condition reported in B. Upregulated genes are indicated in red, and downregulated genes are indicated in green. (D) Venn diagrams comparison of the upregulated and downregulated genes relative to RNAseq conditions reported in A and B; the panel on the right shows a heatmap of the log_2_FC of the shared upregulated (red) and downregulated (green) genes. Predicted allergens are indicated with one asterisk, and two asterisks indicate a predicted secreted lipase.

Next, to elucidate the global transcriptomic response of *M. sympodialis* exposed to nitrosative stress, RNAseq analysis for *M. sympodialis* WT cells treated with sodium nitrite was performed. Compared to the untreated control, 112 genes were upregulated and 50 were downregulated (FDR<0.05; log_2_FC +/-0.5) (Fig. 4B). The most expressed genes included *HEM1* encoding a 5-aminolevulinate synthase involved in heme biosynthesis, *MSYG_3126* encoding a hypothetical secreted lipase, the allantoicase encoding gene *DAL2, MSYG_3153* encoding an uncharacterized NAD(P)/FAD-dependent oxidoreductase, and *DCG1* encoding a protein with unknown function predicted to be related to nitrogen metabolism (Fig. 4B; Dataset S3). The flavohemoglobin-encoding gene *YHB1* was significantly upregulated for FDR <0.05, but it had low expression level (log_2_FC=0.33). Low expression of *YHB1* was also observed in *S. cerevisiae* cells exposed to nitrosative stress (37), although its role in NO consumption has been well characterized (24). The most represented classes of upregulated genes are involved in stress resistance, cellular detoxification and transport, and metabolism (Fig. 4C). Functional protein association network analysis revealed enrichment of genes involved in nitrogen metabolism and regulation, ergosterol biosynthesis, and heat shock response (Fig. S10); we speculate that among the upregulated transcription factor encoding genes (*HSF1, UPC2, BAS1*, and *HMS1*), the heat shock factor Hsf1 might be the key candidate that activates nitrosative stress responsive genes, given its known role in response to stresses in other fungi (38, 39). Conversely, response to nitrosative stress is mediated by the transcription factors Yap1 and Msn2/Msn4 in *S. cerevisiae* and *Schizosaccharomyces pombe* (37, 40), and by the transcription factor Cta4 and the Hog1 kinase in *C. albicans* (41), with the consequent activation of genes known to be required for oxidative stress response, such as those involved in glutathione turnover and other anti-oxidant/detoxification systems. *M. sympodialis CTA1* and *CCP1* are the only oxidative stress responsive genes activated in response to nitrosative stress (Fig. S10; Dataset S3).

The most represented GO category of downregulated genes encodes integral components of membrane, which includes transporters and putative *Malassezia* allergens; other downregulated genes are involved in calcium metabolism, protein folding, and proteolysis. Two transcription factors were downregulated, and they include the pH responsive Rim101, and an uncharacterized bZIP transcription factor (Fig. 4C, Dataset S4).

Comparison of the two different RNAseq datasets revealed two common upregulated genes, encoding the glycerol dehydrogenase Gcy1 and the catalase Cta1, and 10 downregulated genes that include 4 *Malassezia* allergens, a putative secreted lipase, and 5 hypothetical proteins (Fig. 4D). While it is not surprising to find upregulation of a detoxifying enzyme such as catalase, it is intriguing to find downregulation of genes encoding predicted pathogenicity factors.

In conclusion, our transcriptomic data indicate that the response of *Malassezia* to NO and nitrosative stress is mostly different from other studied fungi and it involves metabolic pathways and genes that were not known to be relevant to overcome nitrosative stress.

### *Malassezia* flavohemoglobin has characteristic features of both bacterial and fungal flavohemoglobins

We hypothesized that the structure of a protein acquired by HGT will likely remain similar to that of the donor organism in order to retain its original function. Attempts to resolve the crystal structures of both *Malassezia* flavohemoglobins were carried out, but only the *M. yamatoensis* flavohemoglobin Yhb101 formed crystals to be analyzed. The structure was determined *de novo* by SAD phasing off the heme-iron bound to the globin domain of the protein (Table S3). The flavohemoglobin structure is highly conserved with previously characterized structures of this enzyme family, and it consists of an N-terminal globin domain coordinating an iron-bound (Fe^2+^) heme and a C-terminal reductase domain with both FAD- and NAD-binding sub-domains, of which only FAD is bound (Fig. 5A). An overlay of a flavohemoglobin structure from *E. coli* and *S. cerevisiae* on the *M. yamatoensis* crystal structure highlights conserved binding sites between the proteins (Fig. 5B). Alignment of the globin domains between literature and experimental structures resulted in an RMSD value of 1.532Å for *E. coli* and 1.434Å for *S. cerevisiae*, mostly resulting from slight shifts in the D-loop and E-helix between the structures compared. Common to all structures analyzed is the histidine residue coordinating with the heme iron from the proximal side. This member of the catalytic triad is supported by tyrosine (Tyr98) and glutamate (Glu140) residues conserved in sequence and structure between bacterial, and fungal/yeast flavohemoglobins (42, 43). In *M. yamatoensis*, as also observed in *E. coli*, the heme iron is ligated by 5 atoms: 4 from the heme and His88 from the F-helix. Substrates commonly bind on the distal side of the heme and lead to a conformational change in the planarity of the heme molecule. The E-Helix on the distal side of the heme molecule contributes Leu58, a conserved residue which approaches the heme-bound iron from 3.7Å away. At this position, the 6^th^ coordination site for the iron is occluded, again similar to the *E. coli* crystal structure, but unlike the yeast structure where a three-atom small molecule co-crystallized.

**Figure 5.**
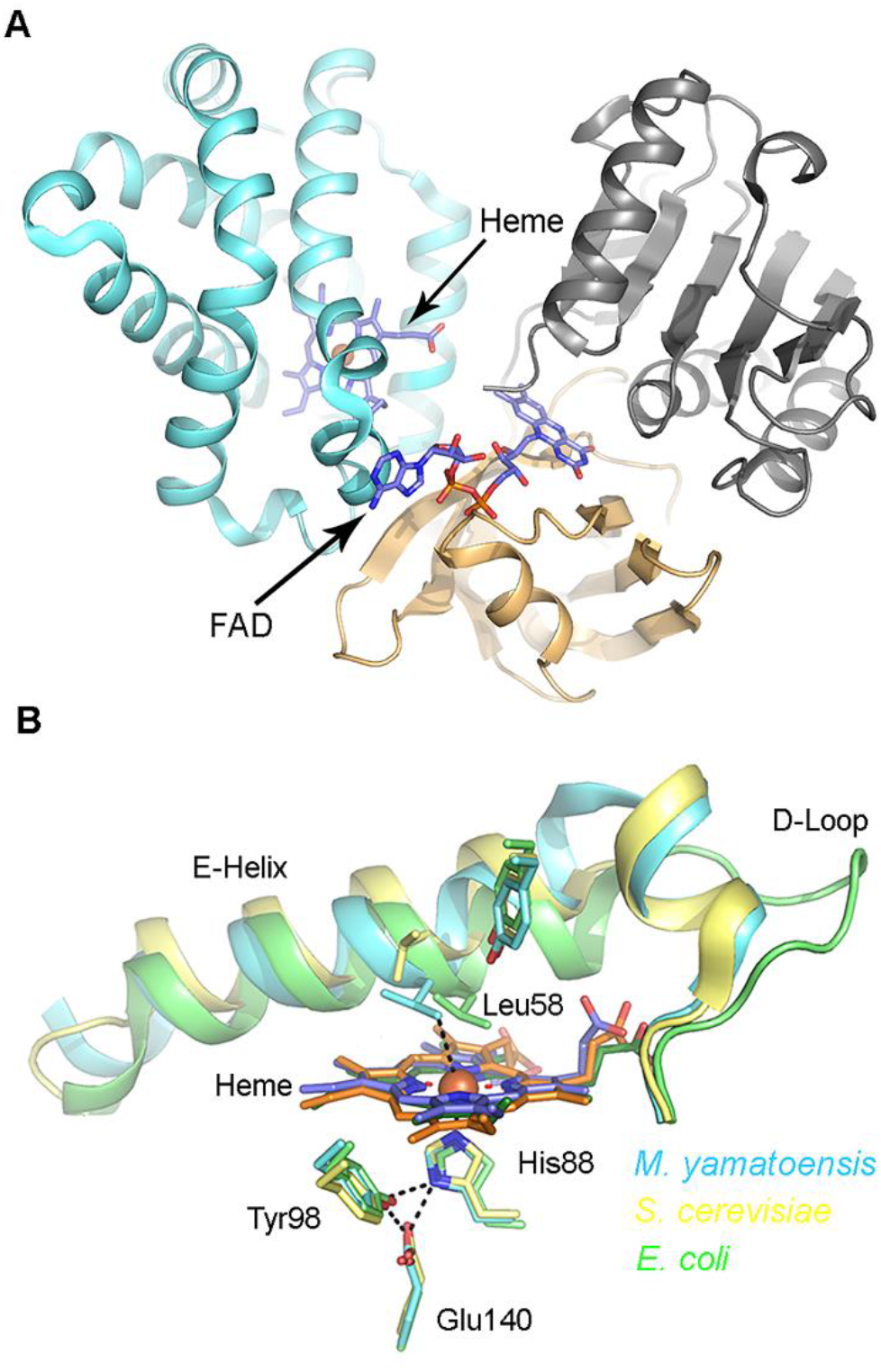
3-dimensional X-ray crystal structure of the *M. yamatoensis* flavohemoglobin Yhb101. (A) The globin domain (cyan) binds a heme molecule. The reductase domain consists of a FAD-binding domain (gray) and a NAD-binding domain (tan) that bind a FAD molecule. (B) An overlay of flavohemoglobin globin domains from fungus, bacteria, and yeast: globin domains of *M. yamatoensis* (PDB ID 6O0A; blue), *S. cerevisiae* (PDB ID 4G1V; yellow), and *E. coli* (PDB ID 1GVH; green) show structural similarity.

In *M. yamatoensis* Yhb101, the D-loop acts as a bridge between the C- and E-helices and the interface between the bound FAD and heme. Comparison of the D-loops from these structures shows the *M. yamatoensis* D-loop adopts a nearly identical helical structure as that of *S. cerevisiae*, in contrast to the *E. coli* D-loop, which is more extended. The *M. yamatoensis* E-helix also adopts a ~30° bend immediately following Leu58, which may straighten out once a substrate is bound. This structural adjustment likely communicates substrate binding near the heme to the reductase domain through movements in the D-loop as the heme B pyrrole proprionate forms a hydrogen bond with the main-chain NH of Ser45, the first residue in the D-loop (Fig. S11).

Lastly, in Fig. S12 a detailed comparison of the functional residues is shown between the *Malassezia* flavohemoglobins with those of the closer HGT donor bacteria *B. ravenspurgense*, *K. kristinae, R nasimurium*, and with the model yeast *S. cerevisiae*.

### *Malassezia* flavohemoglobins are not required for survival on the host

Previous studies in human fungal pathogens indicate that flavohemoglobins are required for pathogenesis (30, 33). In our experiments we found that *M. sympodialis* WT, *yhb1Δ* mutant, and *yhb1Δ* + *YHB1* and *yhb1Δ* + *YHB101* complemented strains have similar levels of survival within activated macrophages (Fig. S13A-B). This result is in contrast with previous findings in *C. neoformans* (30), but it corroborates results obtained in *A. fumigatus* (18). Furthermore, the recently developed murine model for *Malassezia* skin infection (5) was utilized to test pathogenicity of the flavohemoglobin strains and the induction of host response. Corroborating *ex vivo* data, we found no differences both in terms of host tissue colonization and host inflammatory response for the *yhb1*Δ mutant compared to the complemented strains (Fig. S13C-E, Fig. S14). In agreement, there were no differences between WT and *Nos2-/-* mice when challenged with *M. sympodialis* WT (Fig. S13F-H). These results suggest that flavohemoglobin is not required for pathogenesis of *Malassezia* in an experimental skin model.

Lastly, several attempts were also carried out to test survival of *M. sympodialis* WT and flavohemoglobin strains within the GI tract of WT and *Nos2*-/-mice. Because of the high amount of NO produced in the GI tract of mice during inflammation (44), and the recently-reported involvement of *M. restricta* in Crohn disease (6), we hypothesized that the flavohemoglobin would be required for *M. sympodialis* survival in GI tract during inflammation. We followed the protocol developed for GI tract colonization by *M. restricta* (6), but unfortunately we could never recover any *M. sympodialis* colony. For this reason, whether *Malassezia* flavohemoglobin is required for survival within the GI tract could not be determined.

### Analysis of *Malassezia* genomes revealed extensive HGT events from bacteria

Given the gain of function due to acquisition of the bacterially-derived flavohemoglobins by *Malassezia* species, we sought to identify additional HGT candidate genes in the *Malassezia* genus. In a previous study, 8 HGT events were identified in *M. sympodialis*, and then their presence was assessed in other species within the genus (3). In the present study we applied a previously described analytical pipeline (45) based on three HGT metrics - the HGT index (46), the Alien Index (AI) (47) and the Consensus Hit Support (CHS) (48) - to identify novel genus and species-specific HGT events. Our goal was not to explicitly establish the evolutionary history of individual genes, but rather to estimate bacteria-derived HGT candidates for the complete set of *Malassezia* genomes. Besides recovering the *YHB1* and *YHB101* genes as HGT candidates, in addition this analysis identified a total of 30 HGT candidate genes (Fig. 6 and Dataset S5), seven of which in common with the previous study. HGT candidates found in the majority of the *Malassezia* species include genes involved in broad resistance to stresses, including three that were upregulated in *M. sympodialis* exposed to nitrosative stress (Dataset S3), such as the NAD(P)/FAD-dependent oxidoreductase-encoding gene *MSYG_3153*, the catalase-encoding gene *MSYG_3147*, and the sorbitol dehydrogenase-encoding gene *MSYG_0932* (Fig. 6). Other HGT candidates include a deoxyribodipyrimidine photo-lyase predicted to be involved in repair of UV radiation-induced DNA damage, and a class I SAM-dependent methyltransferase potentially modifying a variety of biomolecules, including DNA, proteins and small-molecule secondary metabolites. Another interesting HGT candidate is the gene encoding a septicolysin-like protein, which is known as a pore-forming bacterial toxin that might play a role as virulence factor (49, 50). This gene is absent in all *Malassezia* species phylogenetically related to *M. sympodialis*, and is present as five copies in *M. globosa*. Furthermore, a large number of HGT events unique to *Malassezia* species of clade A were found, and the acquired genes encode a variety of proteins with different functions, such as hydrolysis, protein transport and folding, detoxification of xenobiotics, and resistance to stresses. Finally, 12 of the HGT candidates identified were unique to certain *Malassezia* species. An intriguing case is *M. japonica* for which we found four unique HGT candidates, one of them in 3 copies. These genes encode orthologs of the fungal Gre2 protein, which is known to be involved in responses to a variety of environmental stresses (51, 52).

**Figure 6.**
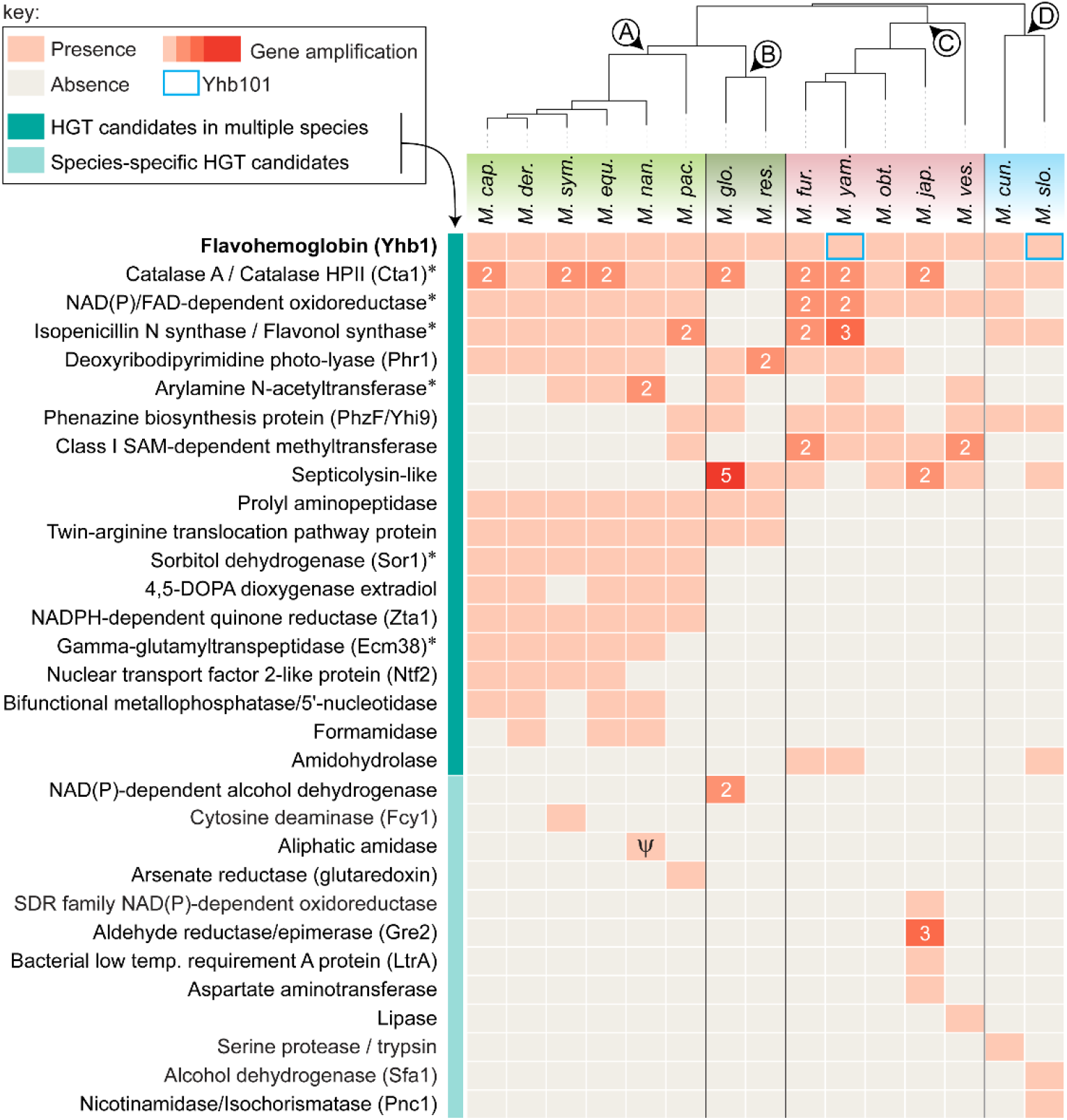
*Malassezia* genes acquired through HGT from bacteria. HGT candidates identified in the genomes of the 15 *Malassezia* species (represented on the top according to their phylogenetic classification) are shown as different lines in the presence-absence matrix, with the closest ortholog in *S. cerevisiae* indicated in parenthesis, where available. For each HGT candidate, the presence of the gene in a genome is indicated by orange square, and the intensity of the color is correlated with the gene copy number (numbers in white). HGT candidates occurring in multiple *Malassezia* species are shown in the top half of the matrix, whereas those that are species-specific HGT candidates are shown in the bottom half of the matrix, and color-coded as shown in the key. Asterisks indicate HGT candidate genes identified in the previous study (3). The bacterially-derived gene encoding an aliphatic amidase identified in *M. nana* CBS9557 seems to be another instance of a pseudogene in this strain (indicated as ψ).

## Discussion

In the present study we report the functional characterization of two *Malassezia* flavohemoglobin encoding genes that were independently acquired through HGT from different Actinobacteria donors. Our experimental analyses demonstrate that both bacterially-derived flavohemoglobins are involved in nitrosative stress resistance and NO degradation, consistent with its known functions in bacteria and fungi (10).

We propose an evolutionary HGT model in which extant flavohemoglobin-encoding genes in *Malassezia* result from a complex pattern of gene retention/loss after being both acquired by a *Malassezia* common ancestor. Nevertheless, other evolutionary scenarios could also be hypothesized, such as: 1) the acquisition of the *YHB1* in a *Malassezia* common ancestor via HGT from a *Brevibacterium-related* donor; 2) followed by more recent acquisitions of *YHB101* by *M. yamatoensis* and *M. slooffiae* via independent HGT events from a common, or closely related, bacterial donor(s) (*Kocuria*). In this scenario, the “resident” *YHB1* in *M. yamatoensis* and *M. slooffiae* could have been displaced upon secondary acquisition of the *YHB101* gene [a phenomenon termed as xenolog gene displacement (53)], or the acquisition of *YHB101* by HGT could have been preceded by the loss of the cognate *YHB1* copy. The identification of novel *Malassezia* species and the analysis of their genomes will be key for the elucidation of these complex models of gene evolution in *Malassezia*.

Although the mechanisms of HGT in fungi are not fully understood, several possible mechanisms have been reported (25, 54). One such mechanism is gene acquisition through conjugation, which requires contact between bacterial donor and fungal recipient (54). For the HGT events that mediated flavohemoglobin acquisition by *Malassezia*, the closest phylogenetic donors are Actinobacteria that are part of the mammalian microbiome and hence share the same ecological niche with *Malassezia*. A dilemma that is common to all HGT events is that if a gene is required for survival in a certain condition, its transfer under that condition might in theory be difficult if not impossible (55). Because NO is synthesized by mammals, including by the skin (56), we speculate that the presence of NO enhanced the HGT transfer of bacterial flavohemoglobins to a fungal *Malassezia* ancestor that acquired the ability to actively consume NO.

Notably, a large number of eukaryotic organisms including fungi lack a flavohemoglobin-encoding gene, suggesting the existence of alternative pathways for nitrosative stress resistance and NO utilization. For example, in *Histoplasma capsulatum*, the etiologic agent of histoplasmosis, Yhb1 is replaced by a P450-type NO reductase (57), whereas in other cases, such as for the basidiomycetous fungi *Moniliella, Ustilago*, and *Puccinia*, NO metabolism in the absence of flavohemoglobin has yet to be elucidated. The evolution and diversification of flavohemoglobin-encoding genes has been a dynamic and complex process characterized by several prokaryote-prokaryote and prokaryote-eukaryote HGT events (22), hence suggesting its significant contribution for habitat colonization by a species but likely a dispensable role in evolutionary divergence.

Is the bacterial flavohemoglobin required for *Malassezia* interaction with the host? While a number of studies in bacteria and fungi reported a role for flavohemoglobin in microbial pathogenesis (10), we surprisingly found that *Malassezia* flavohemoglobins are dispensable for survival within macrophages and for skin infection in our experimental conditions. Conversely, we propose that *Malassezia* flavohemoglobins are important for the commensal lifestyle of *Malassezia* through regulation of NO homeostasis, a hypothesis corroborated by downregulation of genes-encoding putative virulence factors (i.e. allergens and lipases) in our transcriptomic analyses. Another hypothesis is that the HGT-mediated acquisition of flavohemoglobins might be important to mediate *Malassezia* response to NO that is produced by sympatric microbial communities and acts as a quorum signaling molecule, as reported in bacteria (58, 59) and in *S. cerevisiae* (60).

Horizontal gene transfer is thought to occur much less frequently in eukaryotes than in prokaryotes (61, 62), but there are notable cases that invoke HGT as a prominent mechanism of eukaryotic evolution, such as in the transition of green plants from aquatic to terrestrial environments (63), and in the colonization of the animal digestive tracts by rumen fungi and ciliates (55, 64, 65). Analysis of *Malassezia* genomes revealed a large number of HGT events, suggesting that they may also have played a substantial contribution in *Malassezia* evolution and niche adaptation. Donor bacteria include those that are part of the microbiota of animals, but also others that are known to inhabit a variety of terrestrial and marine habitats, raising questions about a possible wider environmental distribution of a *Malassezia* ancestor. This could be correlated with the presence of *Malassezia* DNA in a number of unexpected areas, such as in association with corals and sea sponges in the ocean (66). Moreover, most of the HGT candidate genes identified in *Malassezia* operate as a self-contained metabolic unit, which has been proposed to facilitate HGT (22). Intriguingly, the high number of HGT events suggests also a predisposition of *Malassezia* to bacterial conjugation, in line with our previous findings that *A. tumefaciens*-mediated transformation is the only effective technique for molecular manipulation of *Malassezia* (67). There are a number of identified HGT that are predicted to be important for *Malassezia* pathophysiology and that can be characterized using the methodologies reported in the present study.

## Materials and Methods

### Strains used in the present study

The *M. sympodialis* strains ATCC42132 was utilized as a model species for genetic manipulations. In addition, *M. yamatoensis* CBS9725 (or MY9725) and *M. nana* strains CBS9557, CBS9559, and CBS9560 were employed for NO consumption assays. These strains were grown on modified Dixon’s media (mDixon), which is the medium routinely used for culturing *Malassezia* species (3).

### Identification of *YHB1* genes in *Malassezia* genomes and synteny analyses

Protein-coding sequences were obtained using the *ab initio* gene predictor AUGUSTUS v3.2.2 (68) for each *Malassezia* species that lacked a genome annotation (as of September 2017). To search for *YHB1* homologs, local BLAST databases were set up for both the genome assemblies and the translated coding sequences, and the *M. globosa* Yhb1 protein sequence (GenBank XP_001730006.1) was queried against each database using BLASTP or TBLASTN. Top BLAST-identified protein sequences were retrieved, and the presence of the typical FHbs domains, consisting of an N-terminal globin domain fused with a C-terminal FAD- and NAD-binding oxidoreductase modules, were inspected using InterProScan 5 (69). For the analysis of relative gene-order conservation (synteny) between *Malassezia* species, a region of ~ 32 kb surrounding the *YHB1* gene (~7-8 genes on each side) was carefully examined and compared using top scoring BLASTP against the reference genome annotations of *M. sympodialis*, *M. globosa* and *M. pachydermatis*.

### Yhb1 phylogeny and topology tests

The *Malassezia globosa* Yhb1 protein was queried against the Genbank non-redundant (nr) protein database (last accessed in August 2017) with BLASTP and an e-value inclusion threshold of 1e-10. Protein sequences corresponding to the top 5000 hits were extracted and utilized for downstream analyses. In addition, all putative *Malassezia* Yhb1 proteins identified from the genome data, as well as the functionally characterized Yhb1 gene of *S. cerevisiae*, were included in the dataset. Highly similar sequences were collapsed with CD-HIT v4.7 (70) using a sequence identity threshold of 0.95 (-c 0.95) and word length of 5 (-n 5) to remove redundancy and correcting the bias within the dataset. Sequences were aligned with MAFFT v7.310 using the FFT-NS-i strategy (71) and poorly aligned regions were trimmed with TrimAl v1.4. (-gappyout) (72). The Yhb1 phylogenetic tree was constructed using IQ-TREE v1.5.5 (73) and the LG+F+I+G4 amino acid model of substitution as determined by ModelFinder (74). Because the dataset consisted of a large number of relatively short sequences, we reduced the perturbation strength (-pers 0.2) and increased the number of stop iterations (-nstop 500) during tree search. Branch support was accessed with the ultrafast bootstrap approximation (UFboot) (75) and the SH-like approximate likelihood ratio (SH-aLRT) test (76) both with 10,000 replicates (-bb 10000 –alrt 10000). Tests of monophyly were performed in CONSEL version 1.2 (77) using the approximately unbiased (AU) test (78) to determine whether the maximum likelihood (ML) estimates of the best tree given a constrained topology differed significantly from the unconstrained best ML tree. To produce a constrained topology, *Malassezia* sequences were forced to be monophyletic, and all other branches were resolved to obtain the maximum -log likelihood using RAxML v8.2.11, with five alternative runs on distinct starting trees (-# 5) and the same amino acid model of substitution. Site-wise log-likelihood values were estimated for both trees (option: -f g in RAxML) and the resulting output was analyzed in CONSEL.

### Species phylogeny

To reconstruct the phylogenetic relationship among the 15 *Malassezia* species selected, top pair-wise BLAST results of whole proteomes were clustered by a combination of the bidirectional best-hit (BDBH), COGtriangles (v2.1), and OrthoMCL (v1.4) algorithms implemented in the GET_HOMOLOGUES software package (79), to construct homologous gene families. The proteome of *M. sympodialis* ATCC42132 served as reference and clusters containing inparalogs (i.e. recent paralogs defined as sequences with best hits in its own genome) were excluded. A consensus set of 246 protein sequences was computed out of the intersection of the orthologous gene families obtained by the three clustering algorithms using the perl script *compare_clusters.pl* included in the package. These single copy orthologous gene families were individually aligned with MAFFT v7.310 using the L-INS-i strategy and trimmed with TrimAl (-gappyout). The resulting alignments were concatenated with the python script ElConcatenero (80) to obtain a final supermatrix consisting of a total of 134,437 amino acid sites (47,942 parsimony-informative). The phylogenetic tree was constructed with IQ-TREE v1.5.5 and the LG+F+R5 amino acid model of substitution and branch support values were obtained from 10,000 replicates of both UFBoot and SH-aLRT.

### Horizontal gene transfer analyses

To assess the extent of horizontal transfer into *Malassezia* genomes, we applied a previously described pipeline (45) to the set of 15 available *Malassezia* proteomes, adjusting the parameters accordingly. In brief, protein sequences were first aligned to the UniRef100 database (last accessed September 2017) using Diamond ‘blastp’ (81) with the following parameters: ‘--sensitive --index-chunks 1 --max-target-seqs 500 --Evalue 1e-5’. Three metrics were defined for each protein sequence used as query: (a) the HGT Index (46), (b) the Alien Index (AI) (47), and (c) the Consensus Hit Support (CHS) (48). Hits in Malasseziales were omitted (NCBI taxid 162474), and we specified ‘Fungi’ as the ingroup lineage (NCBI taxid 4751) and ‘non-Fungi’ as outgroup. Proteins receiving an HGT index ≥ 30, AI index > 45, CHS ≥ 90, and that have Bacteria as the donor lineage, were considered well supported HGT candidates and analyzed further. When a given HGT candidate was detected in more than one, but not all the species, TBLASTN searches were used to distinguish between *bona fide* gene losses or genome mis-annotation.

### Molecular manipulation of *M. sympodialis*

A detailed procedure of the cloning procedures and transformation technique used is reported in SI Appendix, SI Material and Methods. The plasmid for targeted mutagenesis of *M. sympodialis YHB1* was generated by cloning regions of 1.5 kb flanking the *YHB1* gene fused with the *NAT* marker within the T-DNA (transfer DNA) of the binary plasmid pGI3 as previously reported (67, 82). For *yhb1*Δ functional complementation, two different GFP-fusion plasmids were generated. The *M. sympodialis YHB1* ORF, the endogenous *M. sympodialis YHB1* gene including its native promoter and terminator, or the endogenous *M. yamatoensis YHB101* gene including its native promoter, were amplified by PCR using chimeric primers that generated recombination sites that allowed T-DNAs assembly in the following order: plasmid pGI35 was *pYHB1, YHB1, GFP, tYHB1, NEO*, and it is referred as *YHB1-GFP;* plasmid pGI31 was *pYHB101, YHB101, GFP, tYHB1, NEO*, and it is referred as *YHB101-GFP*. Plasmids were recombined through *in vivo* recombination in *S. cerevisiae* (67). Correct plasmids were identified by PCR and introduced into the *A. tumefaciens* EHA105 strain by electroporation, and the transformants selected on LB + 50 μg/ml kanamycin.

*M. sympodialis* was transformed through *A.-tumefaciens* mediated transformation following our previously published method (27, 28). Transformants were selected on mDixon supplemented with nourseothricin (100 μg/ml) or neomycin G418 (100 μg/ml), and cefotaxime (350 μg/ml) to inhibit *Agrobacterium* growth. Transformants were purified to single colonies, and subjected to phenol-chloroform-isoamyl alcohol (25:24:1) DNA extraction (83). The correct replacement of the *YHB1* target locus, as well as integration of the reconstitute versions of the *YHB1* and *YHB101* genes, were assessed by PCR. Primer sequences are listed in Table S2.

### NO quantification and GFP expression analysis

Intracellular levels of NO was measured by flow cytometry adapted from previous method (14). In brief, *M. sympodialis* WT and *yhb1*Δ mutant were grown ON in mDixon media, washed once with PBS, and then incubated in PBS with agitation overnight. For each sample, ~ 2 × 10^8^ cells/mL were equally divided, and were stained with DAF-FM DA at a final concentration of 10 μM, and the other half not stained and used as background control. Cells were washed once with PBS and analyzed using a BD FACSCantoTM II.

To measure intracellular GFP intensity by flow cytometry, *M. sympodialis yhb1Δ* mutants, and complemented strains *yhb1Δ* + *YHB1-GFP* and *yhb1Δ* + *YHB101-GFP* were grown in mDixon media ON, washed in PBS, and ~1 × 10^8^ cells/mL used for flow cytometry analysis on the Becton-Dickinson FACScan at Duke Cancer Institute Flow Cytometry Shared Resource. The results were analyzed using FlowJo. *M. sympodialis yhb1Δ* mutants, and complemented strains *yhb1Δ* + *YHB1-GFP* and *yhb1Δ* + *YHB101-GFP* prepared in the same way were also used for GFP microscopy analysis carried out at the Duke Light Microscopy Core Facility using a Zeiss 710 inverted confocal microscope.

### NO consumption assay

The NO consumption assay was performed as previously reported (84, 85). A custom-made glass reaction cell was filled with PBS (4 mL, pH 7.4, 37°C) and connected with a flow meter and a TEA NO analyzer. Under near-vacuum conditions, NO generation was achieved by injecting DETA NONOate (dissolved in 0.01 N NaOH at a concentration of 30 mM) into the reaction cell. Under a helium flow, NO was passed through an inline condenser (removing water vapor) to a NO chemiluminescence analyzer (TEA 810, Ellutia) to generate an electrochemical NO baseline signal (expressed as mV) (Fig. S5).

We established the optimum amount of DETA NONOate to be injected as 30 μL (225 μM), leading to an increase of ~60 mV in the baseline. *Malassezia* yeasts were grown ON in mDixon, washed twice with PBS pH = 7.4, and adjusted in PBS to OD_600_=1. Because *Malassezia* yeasts form clumps in liquid culture, cell counts using a hemocytometer was inaccurate and the number of viable cells in cultures was instead determined by plating an aliquot (100 μL) of the cellular suspensions on mDixon agar and counting colonies. The density of the *Malassezia* cultures was ~2.3 × 10^8^ CFU/mL, with the exception of *M. nana* cultures that were 1.5 × 10^8^ CFU/mL. For NO consumption assays, injection of a volume of PBS pH = 7.4 alone greater than 50 μL interfered with the NO signal baseline, and therefore amounts ranging from 10 μL (2.3 × 10^6^ cells injected) to 50 μL (1.15 × 10^7^ cells injected) of *Malassezia* cellular suspensions were utilized. For *M. nana* injections of 15 μL (2.25 × 10^6^ cells injected), 30 μL (4.50 × 10^6^ cells injected) and 45 μL (6.75 × 10^6^ cells injected) were performed. For technical reasons (i.e. presence of clumps, non-resuspended cells, and cultures too dense for the syringe needle) we found that the indicated range of cell counts were optimal to obtain results that were clean and reproducible.

Our standardized NO consumption assay was performed as follows: once a steady baseline was obtained (approximately 1 min), 30 μL of DETA NONOate was injected, and after a stable NO signal was achieved, different volumes of *Malassezia* cellular suspensions were injected into the buffer solution (PBS + DETA NONOate), resulting in a decrease in NO signal that indicates NO consumption. Following the injection of each *Malassezia* species, the reaction chamber was flushed, and refilled with PBS and DETA NONOate. A maximum of 5 injections was performed using the same buffer solution, after which it became cloudy, generating an unstable signal.

### *In vitro* phenotypic characterization of *Malassezia* strains

Phenotypic analysis of the *Malassezia* strains was performed on mDixon agar by spotting 1.5 μL of 1:10 dilutions of each cellular suspension in the following conditions: DETA NONOate (0.5 mM and 1 mM), NaNO2 (5 mM and 10 mM), hydrogen peroxide (1 mM), UV (200 μJ × 100), 37°C, benomyl (20 μM), 5-flucytosine (5FC, 50 and 100 μg/mL), amphotericin B (AmB, 50 μg/mL), caspofungin (2.5 μg/mL and 5 μg/mL), fluconazole (FLC, 0.5 μg/mL and 1 μg/mL), NaCl (1M), LiCl (100 mM), Congo red (0.4 μg/mL), pH = 7.5, or pH = 4.When hypoxic conditions were required, the GasPak EZ Container System was used (BD Diagnostics). Plates of mDixon supplemented with DETA NONOate (0.1 nM, 1nM, 10 nM and 0.1 mM) and NaNO2 (10 nM, 50 nM, 0.1 mM and 1 mM) were spotted as reported above and placed in the GasPak Large Incubation Chamber with three anaerobe sachets added prior to sealing the chamber. Maintenance of hypoxic conditions (less than 1% O2 and greater than 13% CO_2_) was monitored using Dry Anaerobic Indicator Strips (BD).

### RNAseq analysis

10 mL liquid mDixon cultures of ~ 1×10^7^ cells/mL of *M. sympodialis* WT ATCC42132 with and without 10 mM NaNO2, and *M. sympodialis yhb1Δ* mutant were grown for ON on shaking culture at 30°C. Cells were pelleted at 3,000 rpm in a table top centrifuge for 3 minutes, washed with 10 mL dH2O, and the cell pellet was lyophilized and stored at −80°C until RNA extraction. Cells pellets was broken with sterile beads and RNA extracted with TRIzol (ThermoFisher) according to the manufacturer’s instructions. Final RNA pellets were treated with TURBO DNase (ThermoFisher, catalog # AM2238) according to manufacturer’s instructions, and RNA was resuspended in 50 μl of nuclease free water. Illumina 50 bp single end libraries were prepared using the TruSeq Stranded Total RNA-seq Kit and subjected to Illumina sequencing. Library preparation and RNA sequencing was performed at the Duke University Center for Genomic and Computation Biology. Three biological replicates were performed for each sample.

After sequencing, Illumina raw reads were trimmed with Trimmomatic to remove Illumina adaptors (86) and mapped to the most recent *M. sympodialis* reference genome (87) using HiSat. Generated .bam files were used to run StringTie with the *M. sympodialis* annotation as guide, and the -e option to provide the correct output for the statistical analysis (88). Read count information for statistical analysis were extracted using a provided python script (*prepDE.py*). DESeq2 was used to determine the differentially expressed genes (DEGs) as having FDR<0.05 and log_2_FC ± 0.5, which are common parameters used to define relevant genes in RNAseq experiments (89). StringTie and DEseq2 were run on Galaxy (90). Functional annotation of the DEGs was performed using the Blast2GO pipeline, which includes the BLASTx against the NCBI non-redundant protein database, gene ontology (GO) annotation and InterProScan (91). Venn diagram to identify DEGs in common between the two comparisons were generated using the following web server (http://bioinformatics.psb.ugent.be/webtools/Venn/). Gene network interactions were determined using STRING (https://string-db.org/cgi/input.pl) with the *S. cerevisiae* protein set as reference.

### Flavohemoglobin purification and crystal structure

A detailed procedure for the flavohemoglobin purification and crystal structure is reported in SI Appendix, SI Material and Methods. Briefly, a construct expressing His-Tev-*YHB101* was cloned into *E. coli* BL21(DE3) cells for expression studies. Protein expression was induced with 1 mM Isopropyl β-D-1-thiogalactopyranoside (IPTG), and 5-aminoluevulinic acid 0.3 mM was also added to facilitate heme biosynthesis. The cellular pellet was collected and lysed via microfluidization with two passes at 15,000 PSI on ice and clarified via centrifugation at 200 rcf for 45 minutes at 4°C, and filtered with a 0.2 μm filter. The supernatant was applied to a Ni^2+^ charged HiTrap Chelating HP (GE Healthcare) columns and the protein eluted with a 500 mM imidazole gradient. The fractions of interest (which were visibly red-brown) were pooled, the His-TEV tag removed while dialyzing overnight at 4°C, and fractions of the cleaved protein eluted from the column were concentrated for size exclusion chromatography via centrifugal concentration.

Apo *M. yamatoensis* flavohemoglobin protein was set up for crystallization. Crystals were obtained with Morpheus B12: 12.5% (w/v) PEG1000, 12.5% (w/v) PEG3350, 12.5% (v/v) MPD, 0.03 M each sodium fluoride, sodium bromide, and sodium iodide, and 0.1 M bicine/Trizma base pH 8.5. After 35 days, red-brown crystals were harvested and analyzed on a Rigaku FR-E+ 007 SuperBright rotating anode equipped with Rigaku Varimax optics and a Saturn 944+ detector using several packages described in detail in SI Appendix, SI Material and Methods. The flavohemoglobin structures of *E. coli* (PDB ID 1GVH) and *Saccharomyces cerevisiae* (PDB ID 4G1V) were used for comparison with that of *M. yamatoensis*.

### Interaction of *M. sympodialis* strains with the host

The ability of *M. sympodialis* WT, *yhb1Δ* mutant, and complemented strains *yhb1Δ* + *YHB1-GFP* and *yhb1Δ* + *YHB101-GFP* to survive within macrophage was carried out according to previous protocol (30, 92). J774 A.1 cells (1 × 10^5^/well) were added to 96-well plates and activated by addition of either 1) 10 nM phorbol myristate acetate (PMA) and incubated for 1h at 37 °C with 5% CO_2_, 2) 100 U/mL interferon-γ and LPS (0.6 μg/mL) and incubated overnight at 37°C 5% CO_2_. Two days old cultures of *M. sympodialis* were washed twice with sterile water, and resuspended in RPMI + human serum 20 % for opsonization for 2 h at 37°C. *M. sympodialis* cells were washed three times with water, resuspended in DMEM and added to the macrophages at a 1:1 yeast: macrophage ratio. Plates were incubated for 2 h at 37 °C in 5% CO_2_, then the co-cultures were washed three times with PBS to remove yeasts that were not internalized, and the plate was incubated at 37 °C in 5% CO_2_ for 18 h. Yeast cells were collected by lysing macrophages, and from each condition a 1:20 dilution was plated on mDixon agar and incubated at 30 °C for 3-5 days to determine yeast survival.

For *in vivo* infection experiments, WT C57BL/6j mice were purchased from Janvier Elevage. *Nos2-/-* mice (93) were obtained from Nicolas Fasel (Lausanne). Mice were maintained at the Laboratory Animal Science Center of University of Zurich, Zurich, Switzerland and used at 6-12 weeks in sex- and age-matched groups. Epicutaneous infection of the mouse ear skin was performed as described previously (5, 94). Briefly, *Malassezia* strains were grown for 3-4 days at 30°C, 180 rpm in liquid mDixon medium. Cells were washed in PBS and suspended in native olive oil at a density of 20 ODA_600_/mL. 100 μl suspension (corresponding to 2 ODA_600_) of yeast cells was applied topically onto the dorsal ear skin that was previously barrier-disrupted by mild tape stripping while mice were anaesthetized. For determining the fungal loads in the skin, tissue was transferred in water supplemented with 0.05% Nonidet P40 (AxonLab), homogenized and plated on mDixon agar and incubated at 30°C for 3-4 days. For quantification of cellular infiltrates, single cell suspensions of ear skin were stained with antibodies directed against CD45 (clone 104), CD11b (clone M1/70), Ly6G (clone 1A8), Ly6C (clone HK1.4) in PBS supplemented with 1% FCS, 5 mM EDTA and 0.02% NaN3. LIVE/DEAD Near IR stain (Life Technologies) was used for exclusion of dead cells. Cells were acquired on a FACS Cytoflex (Beckman Coulter) and the data were analyzed with FlowJo software (FlowJo LLC). The gating of the flow cytometric data was performed according to the guidelines for the use of flow cytometry and cell sorting in immunological studies (95), including pre-gating on viable and single cells for analysis. For transcript expression analysis, total RNA was isolated from ear skin according to standard protocols using TRI Reagent (Sigma Aldrich). cDNA was generated by RevertAid reverse transcriptase (Thermo Fisher). Quantitative PCR was performed using SYBR Green (Roche) and a QuantStudio 7 Flex (Life Technologies) instrument for *Il17a* (96), *Defb 3* (97), and *Nos2*. All RT-qPCR assays were performed in duplicates and the relative expression (rel. expr.) of each gene was determined after normalization to *Actb* transcript levels. The primers used for qPCR are listed in Table S2.

### Data Availability

The sequence data generated in this study were submitted to National Center for Biotechnology Information under BioProject accession number PRJNA626605. Individual accession numbers are SRR11574550 for RNA-seq reads of *Malassezia* WT untreated control samples, SRR11574549 for RNA-seq reads of *Malassezia* WT NO-treated samples and SRR11574548 for RNA-seq *Malassezia yhb1Δ* mutant. The final structure factors and coordinates of the flavohemolgobin Yhb101 of *M. yamatoensis* were deposited in the PDB with code 6O0A.

## Supporting information

SI Appendix File_final

## ACKNOWLEDGMENTS

We thank Stephen Rogers for assistance with the NO consumption assay, Nicolas Fasel for *Nos2*-/-mice, Ellen Wallace for her contribution to cloning the *Malassezia* crystallography constructs, Jan Abendroth for his contributions to solving the structure of *Malassezia* by phasing off the bound iron, Jason Yano and Rana Sidhu for consultation on improving heme incorporation during recombinant expression of the crystallography constructs, and Tom Edwards and Don Lorimer for their overall support of the project.

## FUNDING INFORMATION

This work was in part supported by NIH/NIAID R01 grant AI50113-15, and by NIH/NIAID R37 MERIT award AI39115-22 (to J.H.) and by SNF grant 310030_189255 (to S.L.L.). VA Merit BX-003478 supported TJM’s work. Crystallization work was funded by the NIH/NIAID (contract nos. HHSN272200700057, HHSN272201200025C, and HHSN272201700059C to Peter J. Myler). Joseph Heitman is Co-Director and Fellow of the CIFAR program “Fungal Kingdom: Threats & Opportunities”.

## Notes

### Competing Interest Statement

The authors have declared no competing interest.

