## Supplementary material for "Horizontal gene transfer in the human and skin commensal *Malassezia*: a bacterially-derived flavohemoglobin is required for NO resistance and host interaction": SI Appendix File_final

PNAS
www.pnas.org

Supplementary Information for

### Horizontal gene transfer in the human and skin commensal *Malassezia*: a bacterially-derived flavohemoglobin is required for NO resistance and host interaction

Giuseppe Ianiri, Marco A. Coelho, Fiorella Ruchti, Florian Sparber, Timothy J. McMahon, Ci Fu[1], Madison Bolejack, Olivia Donovan, Hayden Smutney, Peter Myler, Fred Dietrich, David Fox III, Salomé LeibundGut-Landmann, and Joseph Heitman*

**This PDF file includes:**

SI Materials and methods

Figures S1 to S14

Tables S1 to S3

SI References

**Other supplementary materials for this manuscript include the following:**

Datasets S1 to S5

#### SI Materials and methods

##### Generation of *M. sympodialis yhb1Δ* targeted mutant and complementing strains through *Agrobacterium tumefaciens*-mediated transformation

For generation of *M. sympodialis yhb1Δ* targeted mutant, 5′ and 3′ ~1.5 kb flanking regions for homologous recombination (HR) were amplified from the genomic DNA of *M. sympodialis* ATCC42132 using primer pairs JOHE44264-JOHE44265 and JOHE44266-JOHE44267, respectively. Primers JOHE44264 and JOHE44267 have chimeric regions for recombination with plasmid pGI3 (1). The *NAT* cassette was amplified from plasmid pAIM1 using primers JOHE43277 and JOHE43278 (2).

For the generation of complementing GFP-fusion strains, the T-DNA cassettes were assembled as follow: the full *YHB1* and *YHB101* genes of *M. sympodialis* and *M. yamatoensis* were amplified from their respective genomic DNA using primers JOHE45236-JOHE45237, and JOHE99961-JOHE99962, respectively. Primers JOHE45236 and JOHE99961 have chimeric regions for recombination with plasmid pGI3. The GFP was amplified from plasmid pSL04 (3) using primers JOHE44555-JOHE45235; primer JOHE44555 has a GGAGGCGGAGGCGGAGGC 6X glycine linker. The *YHB1* terminator was amplified with primers JOHE44557 and JOHE44549; note the same terminator was used for both *YHB1*- and *YHB101*-GFP fusion proteins. The *NEO* marker was amplified from plasmid pAIM5 (2) using primers JOHE44550-JOHE44551; primer JOHE44551 has a chimeric region for recombination with plasmid pGI3. These PCRs were performed using high fidelity (HF) Phusion Taq polymerase (New England Biolabs) using the following conditions: initial denaturation at 98˚C for 30 sec, followed by 32 cycles of denaturation at 98˚C for 20 sec, annealing at 60˚C for 30 sec, and extension at 72˚C for 1 min/kb, and a final extension of 5 min at 72˚C. The PCR products and the double-digested (KpnI and BamHI) binary vector pGI3 were transformed into S. *cerevisiae* using lithium acetate and PEG 3750 as previously reported (2). To assess correct recombination of the newly generated plasmids, single colonies of *S. cerevisiae* transformants were screened by PCR using primers specific for the *NAT* (or *NEO*) marker (JOHE43281–JOHE43282) in combination with primers homologous to outside of the region of the plasmid pGI3 involved in the recombination event (JOHE43279-JOHE43280). Positive clones of *S. cerevisiae* were grown ON in YPD and subjected to phenol-chloroform-isoamyl alcohol

(25:24:1) plasmid extraction using a previously reported protocol (4). These PCRs were carried out using ExTaq polymerase (Taqara Bio, Japan) and the following conditions: initial denaturation at 94˚C for 2 min, followed by 32 cycles of denaturation at 94˚C for 30 sec, annealing at 55˚C for 30 sec, and extension at 72˚C for 1 min/kb, and a final extension of 5 min at 72˚C.The plasmid DNA was then introduced into the *A. tumefaciens* EHA105 strain by electroporation, and transformants were selected on LB + 50 µg/mL kanamycin.

*A. tumefaciens*-mediated transformation of *M. sympodialis* ATCC42132 WT (to generate the *yhb1*Δ mutant) and of *M. sympodialis yhb1*Δ mutant (to generate the complementing strains) was carried out as previously described (5, 6). Briefly, *M. sympodialis* was grown for 2 days at 30°C and the culture was diluted to $OD_{600}$ ~1.0. The engineered *A. tumefaciens* strains were grown overnight, diluted to an $OD_{600}$ ~0.1, and incubated for 4 to 6 h in shaking cultures (30°C) in liquid induction medium (IM) until $OD_{600}$ reached a value of 0.6 to 0.8. The *A. tumefaciens* and *M. sympodialis* cultures were mixed at 1:2 and 1:5 ratios in 50 mL volume, centrifuged at 5200 *g* for 15 min, the supernatants were discarded, and ~500 µl to 1 mL of these fungal and bacterial mixes spotted directly onto nylon membranes placed on mIM agar containing 200 µM acetosyringone. These were coincubated for 7 days at room temperature (plates maintained without Parafilm) prior to transferring the dual cultures to mDixon supplemented with NAT or NEO (100 µg/mL) to select for fungal transformants, and cefotaxime (CEF) (350 µg/mL) to inhibit *Agrobacterium* growth.

*M. sympodialis* transformants resistant to NAT or NEO were colony-purified and subjected to molecular characterization. For the identification of the *M. sympodialis yhb1*Δ mutant, representative NAT resistant colonies were subjected to phenol-chloroform-isoamyl alcohol (25:24:1) DNA extraction as reported above, and analyzed by PCR using primers JOHE44268-JOHE43281 to assess correct junction at the 5′, JOHE44269-JOHE43282 to assess correct junction at the 3′, and with spanning primers JOHE44268-JOHE44269 to assess the correct replacement of the *YHB1* gene with the *NAT* marker through the generation of a larger amplicon in the *yhb1*Δ mutant compared to the WT.

NEO resistant *YHB1-GFP* and *YHB101-GFP* fusion strains were confirmed by PCR using primers JOHE45236-JOHE43278 and JOHE99961-JOHE43278, respectively. The primers used are listed in Table S2.

For RT-qPCR, RNA extraction was performed with the standard TRIzol method, and treated with TURBO DNAse enzyme (Thermo Fisher Scientific). The quality of the RNA was assessed using a NanoDrop spectrophotometer, and 3 µg were converted into cDNA via the Affinity Script QPCR cDNA synthesis kit (Agilent Technologies) according to manufacturer's instructions. A cDNA synthesized without the RT/RNAse block enzyme mixture was used for each sample as a control for genomic DNA contamination. Approximately 500 pg of cDNA were utilized to measure the relative expression level of target genes through quantitative real-time PCR (RT-qPCR) using the Brilliant III ultra-fast SYBR green QPCR mix (Agilent Technologies) in an Applied Biosystems 7500 Real-Time PCR System. Gene expression levels were normalized with the endogenous reference gene *TUB2* and determined using the comparative $\Delta\Delta$Ct method. The primers used are reported in Table S2.

**Flavohemoglobin purification and crystal structure**

The pET28a construct expressing His-Tev-*YHB101* 2-395 (nomenclature relative to SSGCID sub feature MayaA.00765.a) was generated via Gibson assembly as follows. Overlap PCR was used to generate the final PCR product (1305 bp), which was generated from two smaller PCR products: the first PCR product (110 bp) was generated using 20 ng of an unrelated gene as template (amplifying the N-term 8XHis-Tev tag) with primers CID101552_PCR1_For and CID101552_PCR1_Rev, which add a 26 bp 5′ Gibson overlap (homologous with the NcoI-cut end of pET28a) and a 26 bp 3′ overlap with the 5′ end of the second PCR product. The second PCR product (1221 bp) was generated by PCR using 20 ng *M. yamatoensis* DNA as template (amplifying the *YHB101* coding region) with primers CID101552_PCR2_For and CID101552_PCR2_Rev, which add two stop codons and a 30 bp Gibson overlap (homologous with the HindIII-cut end of pET28a) at the 3′ end. The two PCR products were gel-purified on a 1% agarose gel (in 1X TBE + 1X Sybr Green) and 20 ng of each purified PCR was combined and used as template to generate the final PCR using outer flanking primers (CID101552_PCR1_For and CID101552_PCR2_Rev). All PCRs were performed on an ABI Geneamp 9700 using KOD polymerase in the following conditions: initialization at 94°C for 5 min, followed by 28 cycles of denaturation at 94°C for 30 sec, annealing at 65°C for 1 min and extension at 72°C for 1.5 min, followed by a final extension step at 72°C for 7 min.

The resulting amplicon (CID101552_Final) was cloned into the NcoI/HindIII-digested pET28a via Gibson assembly according to manufacturers' instructions, and transformed into *Escherichia coli* TOP10 cells with YT + 50 µg/ml kanamycin for selection. One clone that contained no sequence errors was identified, and the DNA was used to transform *E. coli* BL21(DE3) cells, which served as starting material for expression studies.

Culture of transformed *E. coli* BL21(DE3) was transferred to 2 L of terrific broth containing 50 µg/mL and grown to $OD_{600}$ = 0.6. Protein expression was induced by adding 1 mM Isopropyl β-D-1-thiogalactopyranoside (IPTG) and grown for 45 hours at 25°C. 5-aminoluevulinic acid 0.3 mM was added at the time of induction to facilitate heme biosynthesis. The cells were harvested by centrifugation and the pellets stored at –80°C. Cells were resuspended at 1g:5mL ratios in 25 mM Tris-HCl pH = 8.0 (Teknova), 200 mM NaCl (Teknova), 0.5% 3-[(3-Cholamidopropyl)dimethylammonio]-1-propanesulfonate (CHAPS) (JT Baker), 50 mM L-arginine (Sigma), 500 U of benzonase (Novagen), 100 mg lysozyme (Sigma) and one EDTA free protease inhibitor tablet (Roche). The cells were lysed via microfluidization (Microfluidics) with two passes at 15,000 PSI on ice and clarified via centrifugation at 200 rcf for 45 minutes at 4°C, and filtered with a 0.2 µm PES bottle top filter (Nalgene). The supernatant was applied to four 5 mL $Ni^{2+}$ charged HiTrap Chelating HP (GE Healthcare) columns and the protein eluted with a 500 mM imidazole gradient over 15 column volumes. The fractions of interest (which were visibly red-brown) were pooled and the His-TEV tag was removed via cleavage with His-tagged Tobacco etch virus protease 1 (TEV, produced in-house) while dialyzing against 2 L of 25 mM Tris pH = 8.0 and 200 mM NaCl overnight at 4°C using 10 kDa MWCO snakeskin dialysis tubing (Thermo scientific/Pierce). The affinity tag was removed by applying the digested pool over one 5 mL Ni2+ charged HiTrap Chelating HP columns. The cleaved protein bound and eluted from the column before the remaining uncleaved material and separate pools were made for both. The protein pools were concentrated for size exclusion chromatography via centrifugal concentration (Vivaspin Polyethylsulfone, 10kDa MWCO, Sartorius) to 3.81 mg/mL (cleaved pool) and 14.5 mg/ml (uncleaved pool) for injection over a Superdex75 (GE Healthcare) in 10 mM HEPES (2-[4-(2-hydroxyethyl)piperazin-1-yl]ethanesulfonic acid), pH 7.5 and 150 mM NaCl. Fractions of interest were pooled and concentrated via centrifugal concentration (Vivaspin Polyethylsulfone, 10kDa MWCO, Sartorius) to 5.74 mg/mL (cleaved pool) and 19.93 mg/mL (uncleaved pool), aliquoted and stored at –80°C.

Apo *M. yamatoensis* flavohemoglobin protein (SSGCID ID MayaA.00765.a.TH11.PD38344, uncleaved pool) was set up for crystallization as sitting drops in 96-well XJR trays (Rigaku Reagents) with 0.4 µl reservoir solution and 0.4 µl protein at 19.93 mg/ml. Trays were then stored at 14°C. Crystallization conditions were searched for using commercial sparse matrix screens JCSG+, JCSG-Top96, Wizard 3/4 (Rigaku Reagents), MCSG-1 (Microlytic), and Morpheus (Molecular Dimensions). Crystals were obtained with Morpheus B12: 12.5% (w/v) PEG1000, 12.5% (w/v) PEG3350, 12.5% (v/v) MPD, 0.03 M each sodium fluoride, sodium bromide, and sodium iodide, and 0.1 M bicine/Trizma base pH 8.5. After 35 days, red-brown crystals were harvested directly from the droplet, frozen in liquid nitrogen, and stored for screening.

Data were collected on a Rigaku FR-E+ 007 SuperBright rotating anode equipped with Rigaku Varimax optics and a Saturn 944+ detector. The data set was processed with the XDS package (7). Friedel pairs were kept separate for the calculation of anomalous maps. Initial phases were calculated with Phaser EP (8) within the CCP4 package (9) and the CCP4 program PARROT (10) was used for phase improvement. An initial model was built with ARPwARP (11). The structure was refined with phenix.refine (12) within Phenix (13) and manual model building was done using Coot (14). Built-in tools in Coot were used to assess structure quality before each refinement. Ligand restraints for bound heme and FAD were generated using Grade web server (http://grade.globalphasing.org) from GlobalPhasing Ltd. Model quality was validated using MolProbity (15). The final structure factors and coordinates were deposited in the PDB (16, 17) with code 6O0A. Structural images were generated using Chimera software (18). The flavohemoglobin structures of *E. coli* (PDB ID 1GVH) and *Saccharomyces cerevisiae* (PDB ID 4G1V) were used for comparison with that of *M. yamatoensis.*

#### Supplementary figures

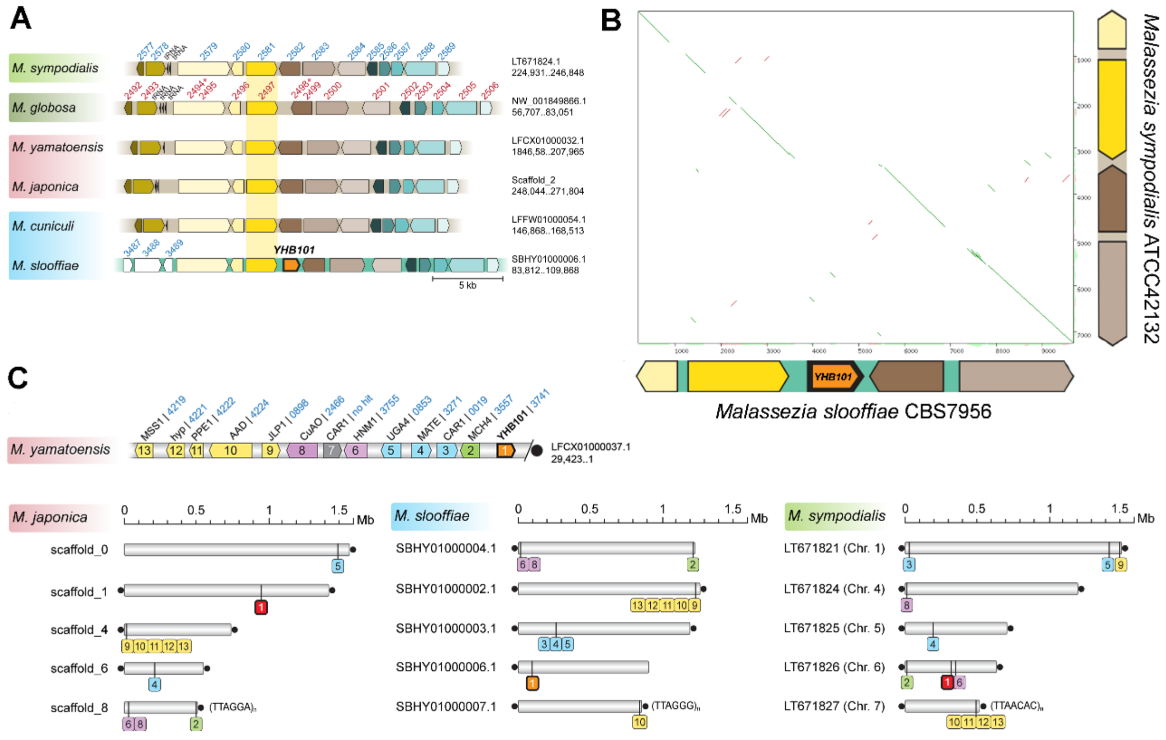

**Fig. S1.** Genomic analysis of the *YHB101* gene in *M. yamatoensis* and *M. slooffiae*. (A) Genomic location of the *YHB101* gene in *M. slooffiae* and comparison with the homologous region in representative species of other *Malassezia* clades. Synteny is well conserved between all species, except for the presence of *YHB101* gene in *M. slooffiae*, as confirmed by the dot plot comparison reported in B. (C) Genomic context of the *M. yamatoensis YHB101* showing its location near the telomere and absence of synteny across species. Orthologs of the *YHB101* and color-coded flanking genes of *M. yamatoensis* are represented for three representative *Malassezia* species highlighting extensive gene reshuffling. Black circle represents the end of a chromosome; scaffold/chromosomal locations and accession numbers are given for each region.

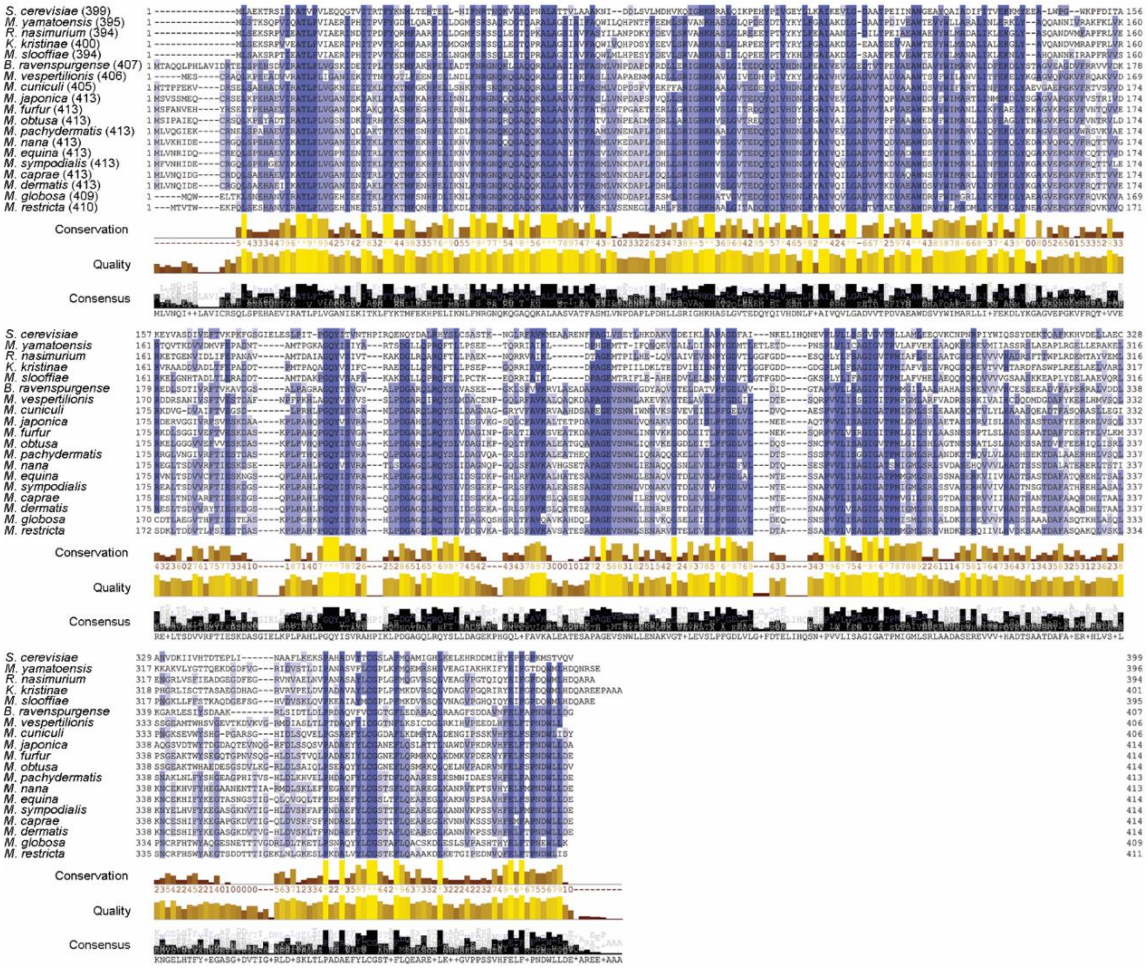

**Fig. S2.** Alignment of the flavohemoglobin Yhb1 of the 15 *Malassezia* species and the predicted donor bacteria *B. ravenspungense*, and of the flavohemoglobin Yhb101 of *M. yamatoensis and M. slooffiae* and the predicted donor bacteria *R. nasimurium* and *K. kristinae*; *S. cerevisiae* Yhb1 is included as a reference. In parenthesis is indicated the protein length. Sequences were aligned with OMEGA and visualized with Jalview (http://www.jalview.org/) using the percentage of identity option: dark blue indicate >80% identity with the consensus sequence, while white is <40% identity with the consensus sequence. Conservation shows a quantitative numerical index reflecting the conservation of the physico-chemical properties for each amino acid, and it is representing the alignments as histogram giving the score for each column. Amino acids with substitutions in the same physico-chemical class have the next highest score. Conserved residues are indicated by * (score of 11 with default amino acid property grouping), and residues with substitutions where all properties are conserved are marked with a + (score of 10, indicating all properties are conserved). Alignment quality annotation is an ad-hoc measure of the likelihood of observing the substitutions in a particular column of the alignment: higher histograms represent conserved residues. Alignment consensus annotation reflects the percentage of the different residue per column and it is overlaid with a sequence logo that reflects the symbol distribution at each column of the alignment.

A

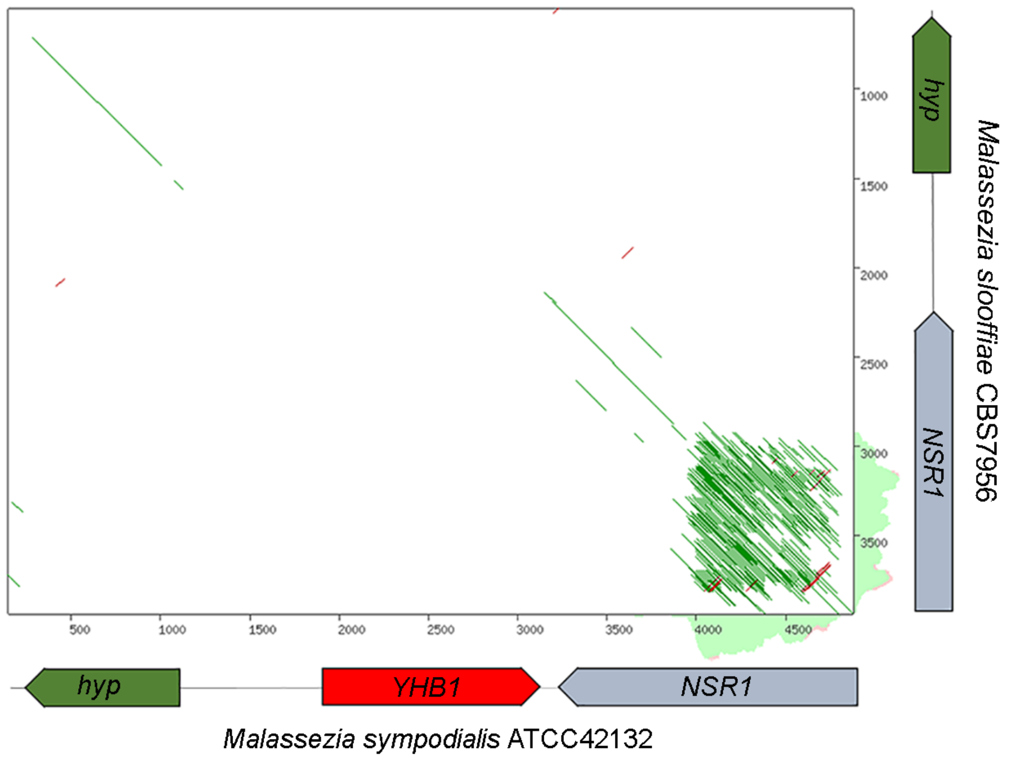

B

>MSYG_3470-*NSR1*
ATGGGCAAGAAAGATAAAATTGCAGCTGCCCCTGTCAAGGCATCGAAGAACGTAAAGGCCTCTACCCGCAAGGGAGA
GACTGCCACTGAAAAGAAGAAGGCGGTGAAGGCGAAAGAGCCTGAACCTATGGAGGAGGAATCTGAAGATGAGCCGG
AGGACTCGGACTCCTCAGAAGACTCTTCGGATGAATCCTCAGACAATTCGTCGAGC**GATGATGACGATTCCTCGGAC**
**GATTCTTCAGACAATGAATCCGATGCTGATTCTAACGATTCCGATGAAGACTCTGATGAGGACGACGACGAGGAGAA**
**AAAGAACTCAAACGACGACTCTGATGATTCCTCAGACGATGATTCTGACGATGATTCTGACGATGACTCCGACGAGG**
**ACGACAATGATGAAAAGAGCTCGAAAGAAGACTCTAGCGATTCCTCTGACGATTCCTCAGATGATTCTTCTGACGAC**
**TCCGATGAGGATAACGATGAAGGCAAGAAGTCAAAAGAAGAAGACTCAGACAACGACTCGGACGACGATTCTGATGA**
**CGATTCTGACGATGACTCGGACGACGACTCTGACTCTGACGACGATTCTGACGATGACGACGATAAAGACAAGAAGA**
**CGTCGAAGAAAGATTCGGATGATGACTCGGATGACGACTCGGATGATGACTCGGATGATGACTCGGATGATGACTCG**
**GATGATGATTCGGATGATGACTCCGACGACGATAAAGACAAGAAGACGTCGAAGAAAGATTCGGATGATGACTCGGA**
**TGACGACTCGGATGACGACTCGGATGACGACTCGGATGACGACTCGGATGACGACTCGGATGACGACTCGGATGACG**
**ACTCGGATGAAGACTCGGACGACGAC**TCGGATGACTCAGGTGAATCCAATGATGACTCGAGCTCGTCTGAAGACGCT
CCCAAGCCCGTATCGAAGAAGCGCAAGGCTGATGACGGTGAAGAAGCCGCACCGAAGAAGACTAAAGTTGACGAGAG
TGTTGACGAAGGTATCAAGACTCTGTGGGTCGGCCAGTTAAGCTGGAACGTTGACAATGACTGGCTGAAAAGTATAT
TCGAAGAGTACGGCACCGTGACGGATGCGCGTGTACAGTGTGACCGCGACTCTGGTCGTAGCCGGGGCTTTGGCTAC
GTCGACTTTGCTACGTCTGCTGAGGCTCTGGAGGCATCCAAGAAGGCGCAGGGCAAAGAGGTCGACGGTCGCAACCT
TCGCGTTGATTTGCAGGCTCCTCGCGCTCCCAAGGAGCGTGCCGATTCGCGTGCCAAGCAGTTTAACGATGAGCGCA
GTGCTCCCTCTAACACACTCTTCCTTGGTGGTCTCGCATGGTCTTTGACTGAAGACGATATCTGGAATACCTTTGCT
GAGTACGGTGAAGTGTCTGCCGTCCGTCTGCCAAAAGAGATTGATTCAGGCCGTCCCAAGGGCTTCGGCTATGTTGA
ATTCGCTTCTCAGGAAAATGCTGCGCAGGCTCTCGAGGCTCTCCATGGACAGGAGCTCGGCGGCCGTCCTATCCGCA
TTGACTTTGCGGGCAAGCGCGACAACACTAACTCTCCACGTGGCGGCGCCTCCAACCGAGGCCGAGGAGGATCCCGC
GGCGGAGCTCGTGGCGGCCGTGACAACGGCTGGGGATCTCGTGCCTCTGGCAGTGCTCGCAGCGGTGCAATCGCGAA
GTCTGCCGGTAAAAAAATGACATTCGACGACTAA

**Fig. S3.** The presence of a repetitive region might have facilitated the flavohemoglobin *YHB1*-mediated HGT event. (A) Pairwise alignment of a 5-kb region surrounding the *M. sympodialis YHB1* gene with the corresponding region in *M. slooffiae*; arrows represent the genes in the selected regions. Green lines indicate forward alignment, while red lines indicate reverse alignment. Note in *NSR1* gene the repetitive region indicated by the green and red lines off the main green diagonal. Alignment was visualized with YASS (https://bioinfo.lifl.fr/yass/yass.php). (B) Sequence of the *M. sympodialis NSR1* gene with the repetitive region represented in bold.

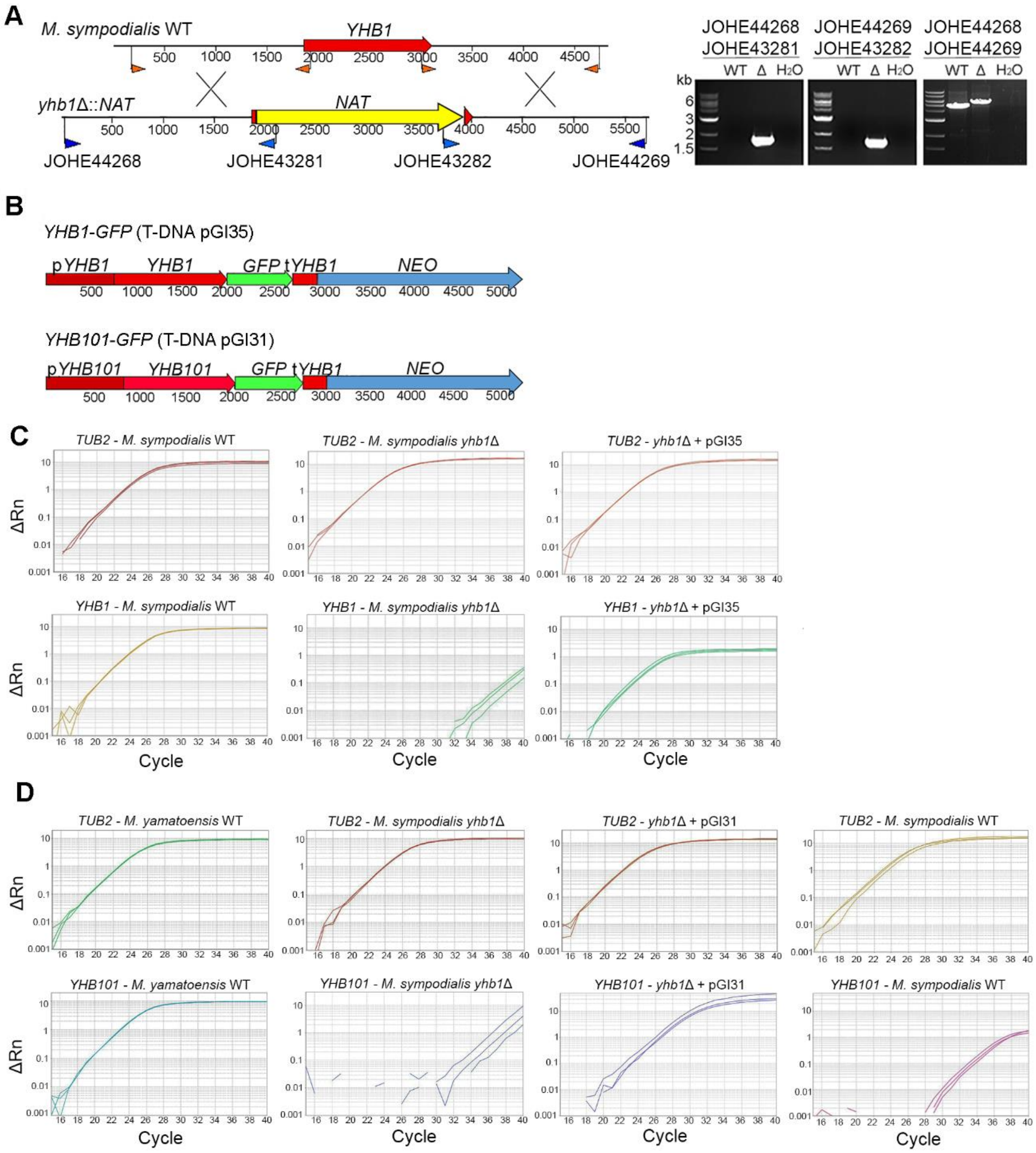

**Fig. S4.** (A) Schematic representation of the targeted gene replacement strategy used to generate *M. sympodialis yhb1*Δ mutants. In the top panel, the large red arrow represents the *M. sympodialis YHB1* gene, and the small red arrows represent the primers used to amplify flanking regions for homologous recombination (see Table S2). In the lower panel is reported the mutated *yhb1*Δ gene, with the large yellow arrow representing the *NAT* marker, and the small blue arrows representing primers used to detect homologous recombination events through PCR as reported in the agarose gel. (B) Schematic representation of the T-DNA of plasmids pGI35 (p*YHB1*, *YHB1*, GFP, t*YHB1*, *NEO*) and pGI31 (p*YHB101*, *YHB101*, GFP, t*YHB1*, *NEO*). (C) RT-qPCR amplification curves relative to *M. sympodialis* WT, *M. sympodialis yhb1*Δ, and complementing strain *M. sympodialis yhb1*Δ + pGI35. In the top panels is shown the housekeeping gene *TUB2*, while in the lower panels is shown the flavohemoglobin encoding gene *YHB1*. (D) RT-qPCR amplification curves relative to *M. yamatoensis* WT, *M. sympodialis yhb1*Δ, complementing strain *M. sympodialis yhb1*Δ + pGI31, and *M. sympodialis* WT. In the top panels is shown the housekeeping gene *TUB2*, while in the lower panels is shown the flavohemoglobin encoding gene *YHB101*. *M. sympodialis* WT was used as control to test the specificity of the primers for the *YHB101* gene of *M. yamatoensis.* In both C and D, in the y axes are reported ΔRn, which represent the magnitude of the fluorescence signal generated during the PCR at each time point; in the x axes are reported the PCR cycles.

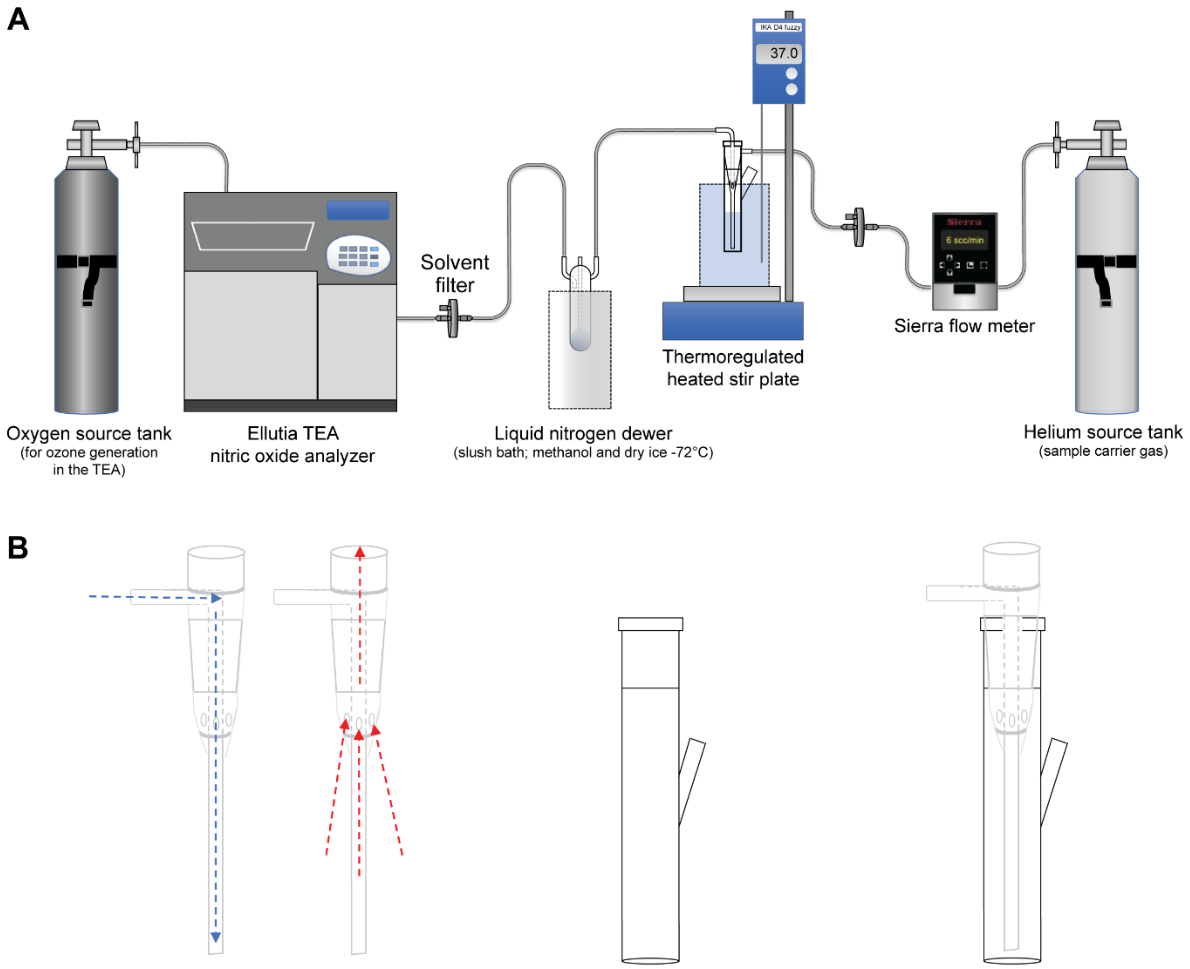

**Fig. S5.** Biochemical assay to detect NO consumption of *Malassezia* strains used in this study. (A) Experimental set up for the biochemical NO scavenging assay. (B) Details of the custom-made glass reaction cell showing gas inflow (blue) and outflow (red) from the chimney (left side), the empty reaction cell (middle), and the chimney and reaction cell fully assembled (right side).

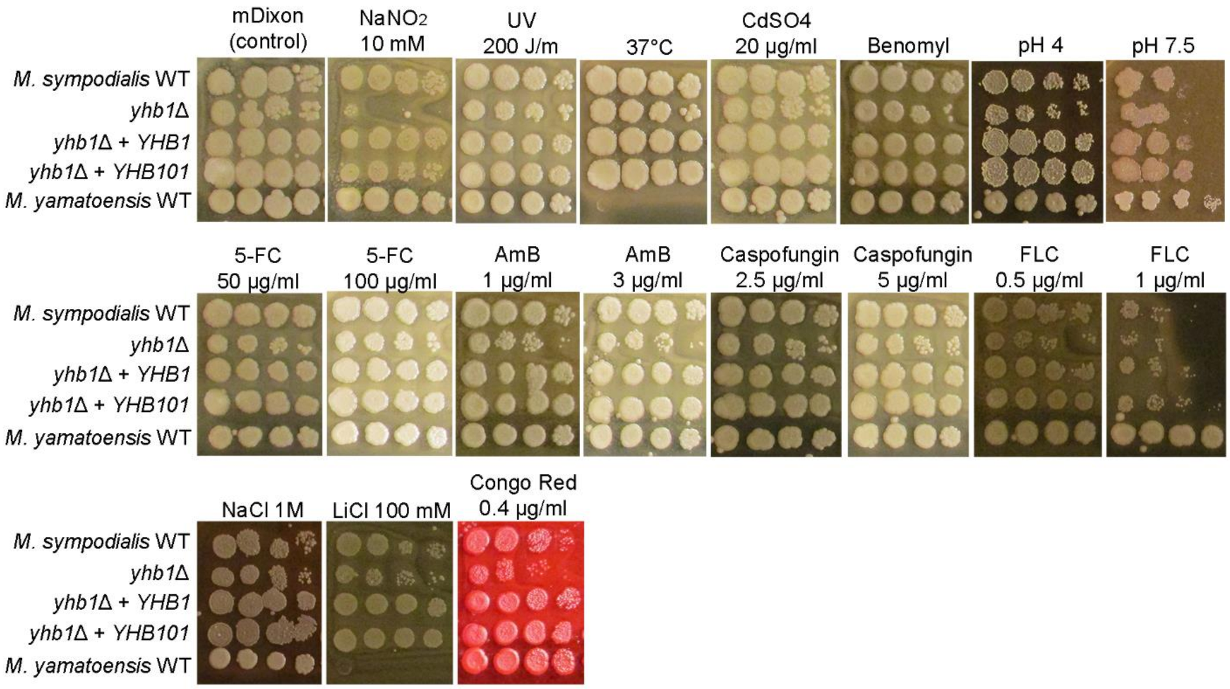

**Fig. S6.** Phenotypic analysis of *M. sympodialis* WT, *yhb1*Δ, *yhb1*Δ + *YHB1*, *yhb1*Δ + *YHB101*, and *M. yamatoensis* WT. Tenfold serial dilutions of cellular suspensions of each strain were spotted (1.5 µL) on mDixon agar containing the indicated stressors.

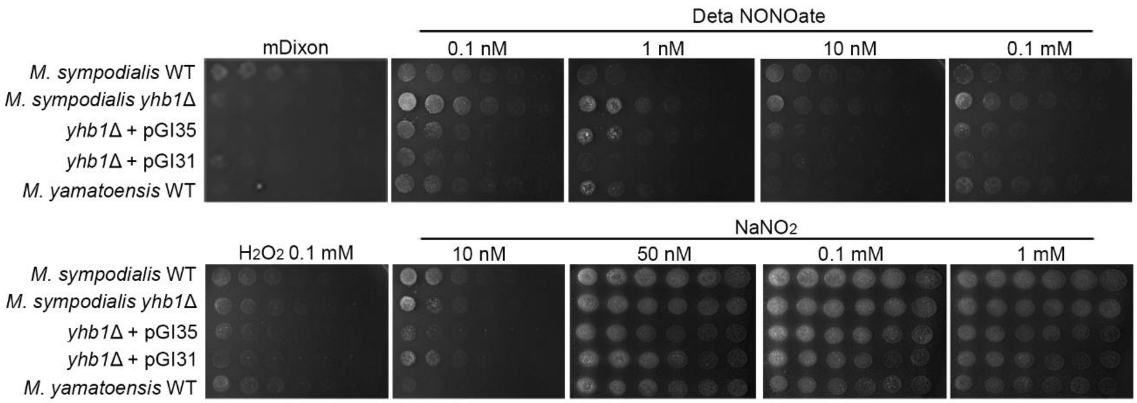

**Fig. S7.** Nitrosative stress resistance analysis of *M. sympodialis* WT, *yhb1*Δ, *yhb1*Δ + *YHB1*, *yhb1*Δ + *YHB101*, and *M. yamatoensis* WT under anaerobic conditions. Tenfold serial dilutions of cellular suspensions of each strain were spotted (1.5 µL) on mDixon agar containing the indicated stressors and incubated for 4-10 days in an anaerobic chamber.

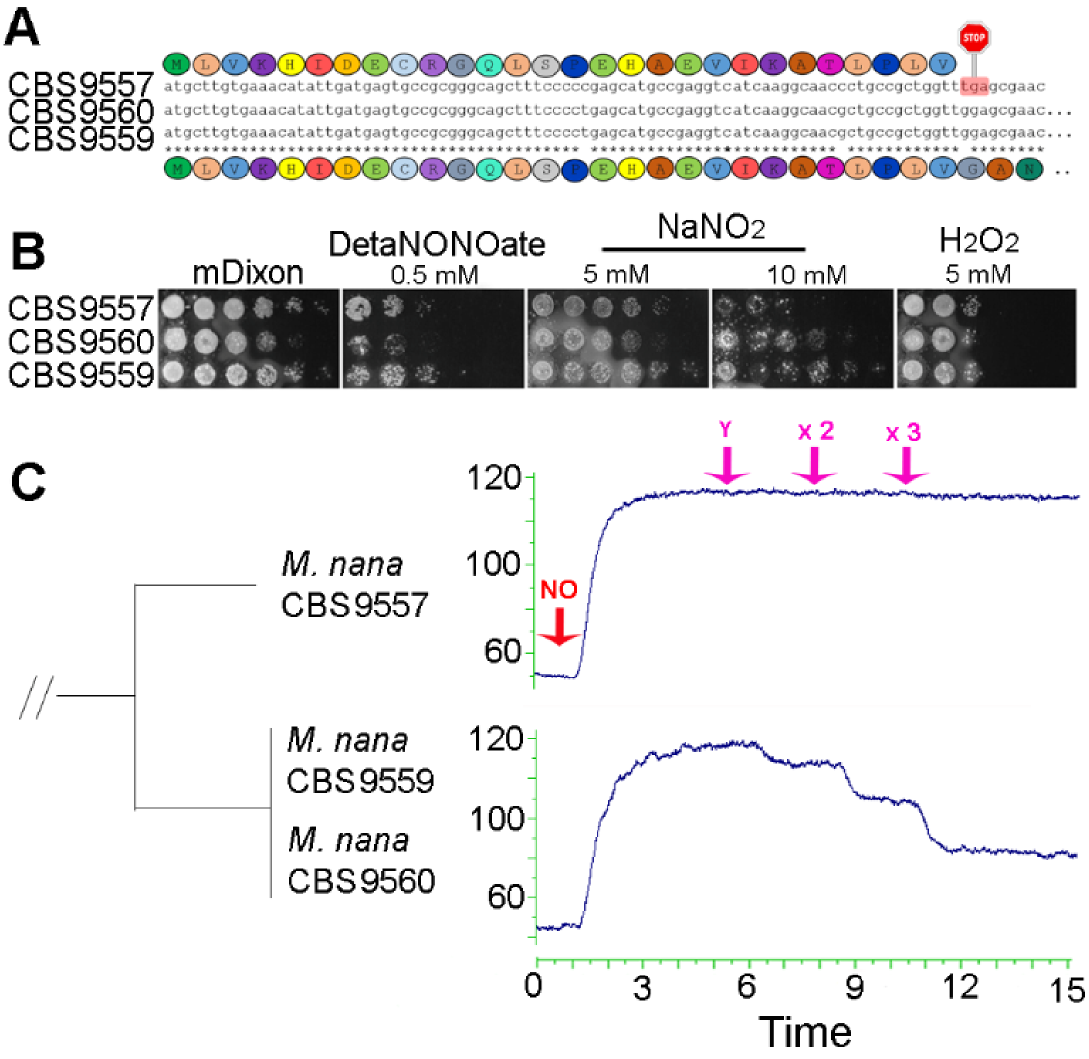

**Fig. S8.** (A) Alignment of the initial ~100 nucleotides of the gene *YHB1* in 3 representative *M. nana* species; on the top of the alignment is reported the protein translation for *M. nana* CBS9557, on the bottom the protein translation for *M. nana* CBS9560 and CBS9559. Each amino acid is depicted in different colors, and the stop sign in *M. nana* CBS9557 indicates the presence of a stop codon. (B) Stress sensitivity assay of *M. nana* CBS9557, CBS9560 and CBS9559 on mDixon agar supplemented with the NO-donor agent DETA NONOate and $NaNO_2$, and with hydrogen peroxide. (C) For the NO consumption assay, 15 µL (Y), 30 µL (x 2) and 45 µL (x 3) of *M. nana* cellular suspensions were injected. The two graphs of NO consumption correspond to *M. nana* geographically-distinct isolates as represented by the phylogenetic tree [from (19)].

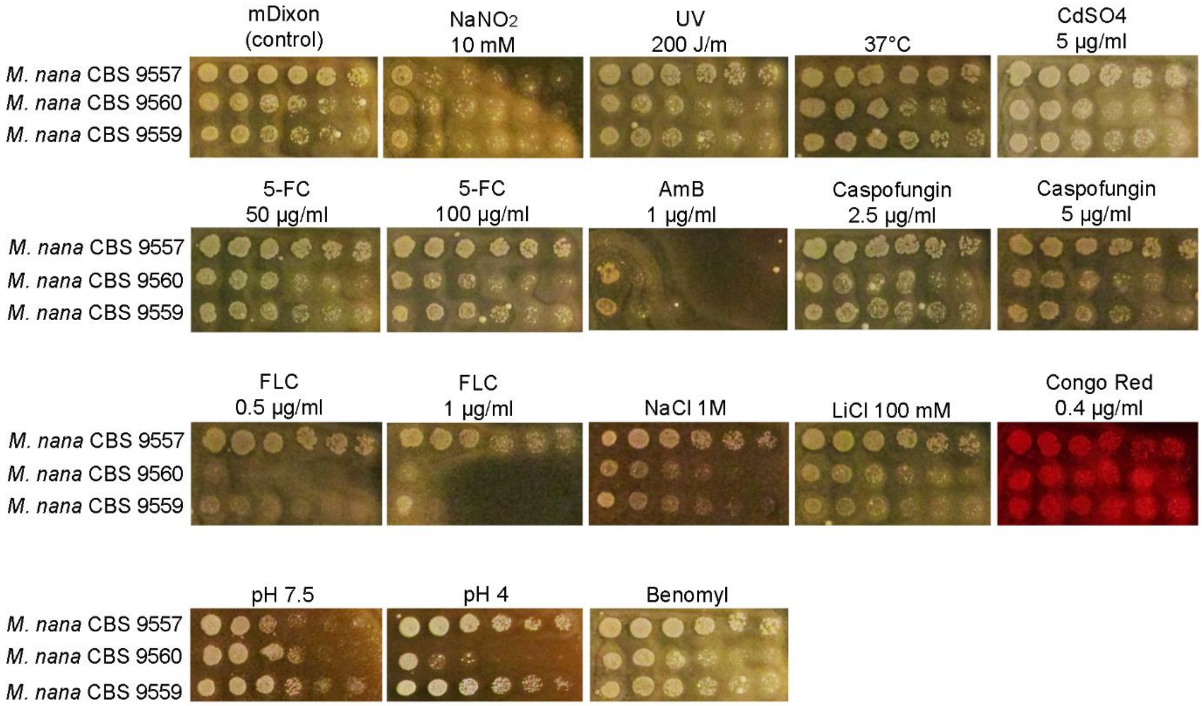

**Fig. S9.** Phenotypic analysis of *M. nana* WT strains CBS9557, CBS9560, and CBS9559. Tenfold serial dilutions of cellular suspensions of each strain were spotted (1.5 µL) on mDixon agar containing the indicated stressors.

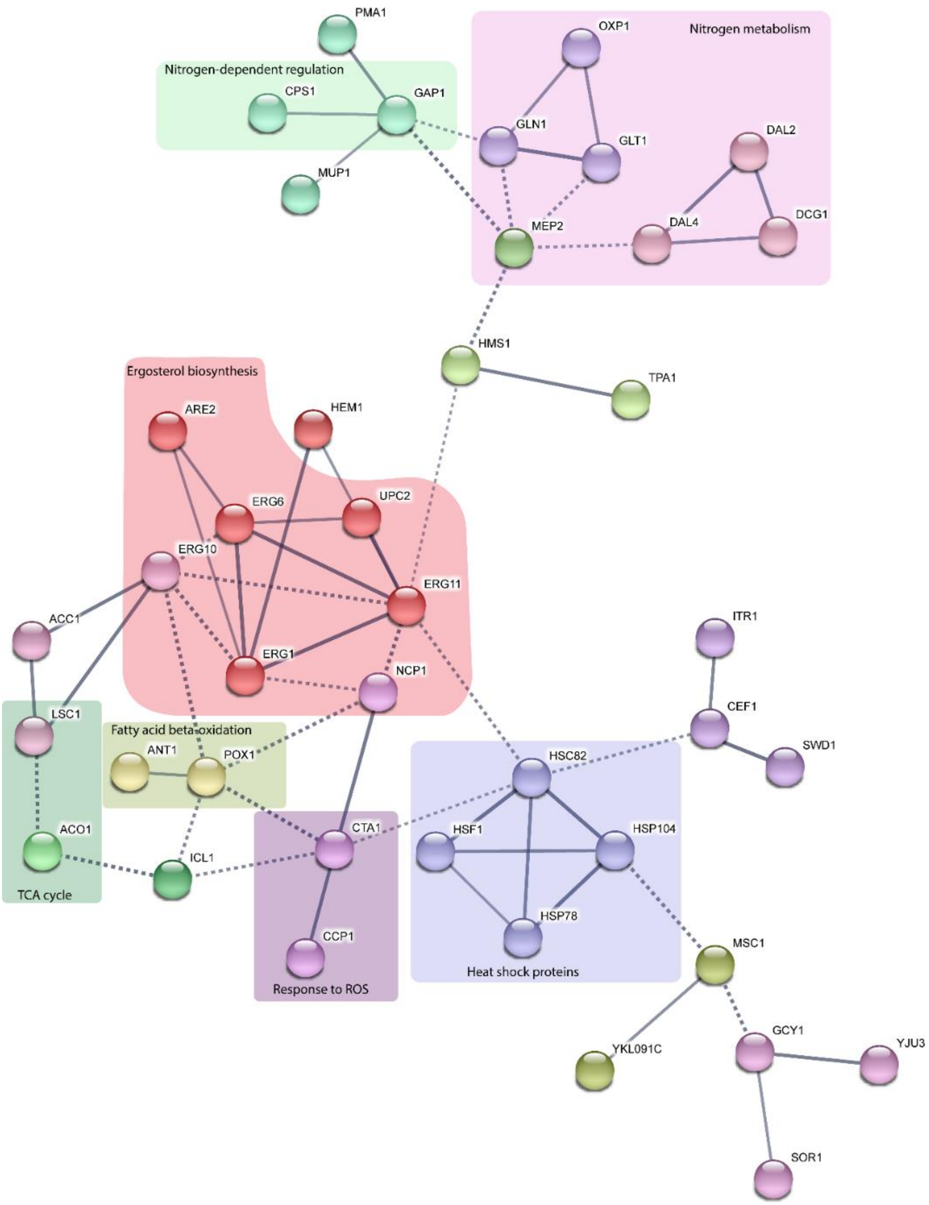

**Fig. S10.** Functional protein association network of upregulated genes of *M. sympodialis* WT strain ATCC42132 in response to nitrosative stress generated by sodium nitrite. Protein names correspond to *S. cerevisiae* orthologs. The network was generated using STRING (https://string-db.org/) with parameters set to draw high confidence interactions with a minimum score of 0.7 (the confidence score is the approximate probability that a predicted link exists between two enzymes in the same metabolic map in the KEGG database); line thickness indicates the strength of data support. Proteins were clustered using the MCL clustering algorithm using an "inflation" value of 3 (inflation is indirectly related with the precision of the clustering, i.e. higher is the inflation more clusters are obtained); every color corresponds to a cluster, and inter-cluster edges are represented by dashed-lines. Based on the role of the identified protein in fungi, known metabolic routes were manually highlighted.

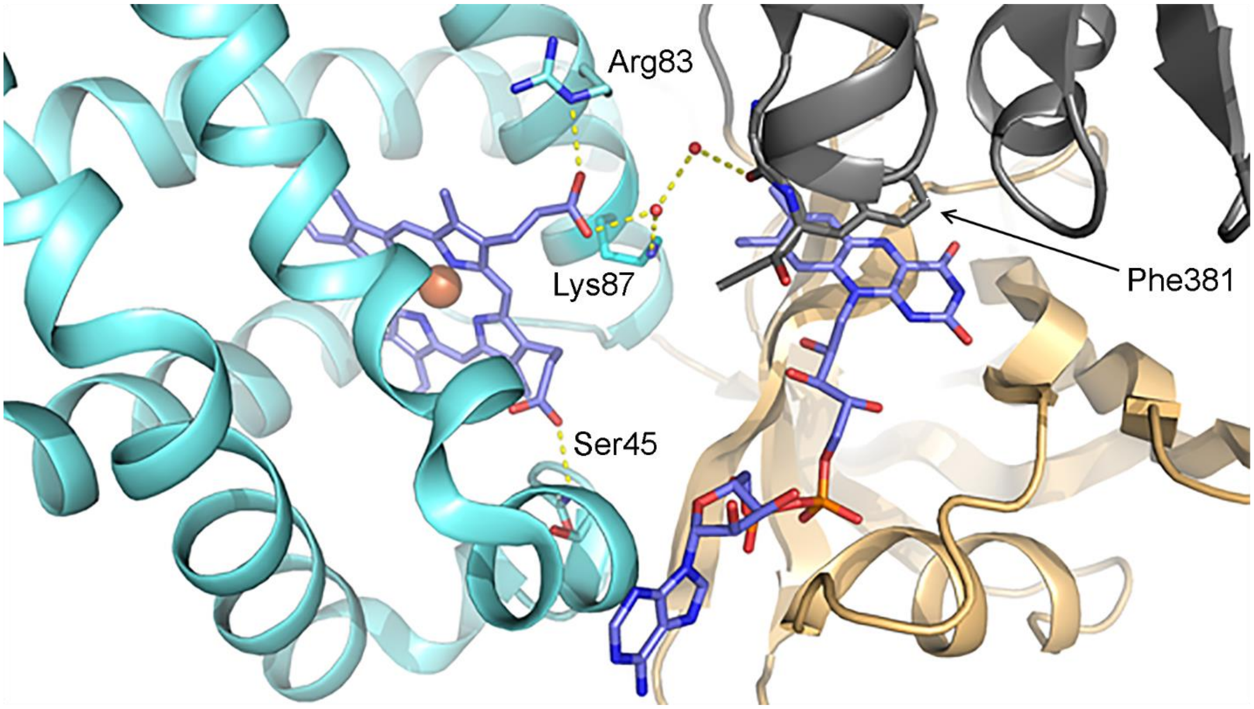

**Fig. S11.** *M. yamatoensis* flavohemolgobin sidechain interactions with heme and FAD. Globin domain and FAD-binding domain directly interact with bound ligands. Residues Ser45, Arg83, and Lys87 bind the E- and F-helices to the heme molecule. Residues Phe381 of the reductase domain and Lys87 of the globin domain, through an interaction with a water molecule, stabilize FAD within the FAD binding domain.

```
S_cerevisiae        ---------------MLAEKTRSIIKATVPVLEQQGTVITRTFYKNMLTEHTELL-NIFN  44
M_yamatoensis       ---------------MLSTKSQPVIQATLPVIAERIPHITPVFYGDMLQARPDLLDGMFS  45
R_nasimurium        ---------------MLSEKSRPVIEATLPVIAERINDITPDFYQRMFAARPDLLDGMFS  45
K_kristinae         ---------------MLSEKSRPVVEATLPVIAERIPHITPVFYRDMFAARPDLMDGMFS  45
M_slooffiae         ---------------MLSAKSRPTIEATLPVIAERIPHITPVFYKDMFDERPDLLNGMFS  45
B_ravenspurgense    MTAQQLPHLAVIDRTELSPEHSDVIRATLPLVGSKIDEITPLFYSKMFAKHPELIADTFN  60
M_vespertilionis    ---------MESCRAQLKPEEADVVRATLPLIGANIEKITTNFYGTLFEENPSLLNDLFN  51
M_cuniculi          ----MTTPFEKVDRSELSAEHADVIRATLPLVGKNIEEITKLFYKTMFGKHPELLSNKFN  56
M_japonica          ----MSVSSMEQCRSELKPEHAEVIRATLPLVGANIEKIAKLFYATMFKAHPELINNLFN  56
M_furfur            ----MSFANVEHYRSELTPEHAEVIRATLPLVGANIDKIAKQFYASMFEGHPELIRNLFN  56
M_obtusa            ----MSIPAIEQCRSQLKPEYADTIRATLPLVGSNIDKITRHFYKSMFEAHPELINNMFN  56
M_pachydermatis     ----MLVQGIEKCRNELSPAHAEVIRATLPLVGANIQDIAKTFYKTMFSNHPELIKDLFN  56
M_nana_CBS9560      ----MLVKHIDECRGQLSPEHAEVIKATLPLVGANIEKITKLFYKTMFEKHPDLIKNVFN  56
M_equina            ----MLVNRIDQCREQLSPEHAEVIKATLPLVGPNIENITRLFYKTMFEKHPELIKNVFN  56
M_sympodialis       ----MFVNHIDECRGQLSPEHGEVIKATLPLVGANIENITKLFYKTMFEKHPELIRNVFN  56
M_caprae            ----MLVNQIDGCRDQLSAEHAEVIKATLPLVGANIENITRLFYQTMFEKHPELIKNVFN  56
M_dermatis          ----MLVNQIDECRGQLSAEHAEIIKATLPLVGANIENIAKLFYKTMFENHPELIKNLFN  56
M_globosa           ---------MQWELTKLSNEHANVIRATLPLVGEHIEEITKLFYKTMFSNHPDLIKNLFN  51
M_restricta         -------MTVTWEKPQLSESHANVIRATLPLVGEHINEITSLFYKTMFQNHPDLIKDLFN  53
                                    *       :.**:*::  .   *:  **  ::  . .*: . *.

S_cerevisiae        RTNQKVGAQPNALATTVLAAAKNI--DDLSVLMDHVKQIGHKHRALQIKPEHYPIVGEYL  102
M_yamatoensis       RSAQRDGTQARALAGSIAIFAQWILQHPNTFPEEMLSRVANKHASLGLQPDEYDTVYKYL  105
R_nasimurium        RSSQLEGTQPKALAGSIAVFASYILANPDKYPDEVLSRVAHKHASLGLREEEYPTVYKYL  105
K_kristinae         RSSQLEGTQPRALAGSIAVFAQWIVQHPDSYPEEVLSRVAHKHASLGLRPEEYPTVYRYL  105
M_slooffiae         RSSQLEGTQPRALAGSIAVFAQWLMLHPESYPDEVLSRVAHKHASIGLQPEEYPTVYKHL  105
B_ravenspurgense    RGNQKLGAQQRALAASIATFASMLVNPDAPDPRDLLERIGHKHASLGITEDQYQIVHDNL  120
M_vespertilionis    RGNQKQGAQQRALAGSIAKFASLLVAPAENMPADLLSRIGHKHAVLGIVEDHYPIVYKYL  111
M_cuniculi          RSNQKQGAQQRALASSIATFASMLVDPDSPVPEKFFARIGHKHAALGVTEDEYQIVHDNL  116
M_japonica          RGNQQQGAQQKALAASVATFASLLVNPDAPSPVQLLSRIGHKHVSLGVTEDQYQIVHDNL  116
M_furfur            RGNQKQGAQQRALAASVATFATMLVTPGAETPERLLGRIGHKHVSLGITRDQYQIVHDYL  116
M_obtusa            RGNQRQGAQQRALAASVATFATVLVNPEADMPDRLLARIGHKHVSLGVTREQYQIVHDYL  116
M_pachydermatis     RGNQKQGAQQKALAASVATFAAMLVNENAPLPDHLLSRIGHKHAALGVTEDQYQIVHDNL  116
M_nana_CBS9560      RGNQQQGAQQKALAASVATFASMLVKEDAPLPDHLLSRIGHKHVSLGVTEDQYQIVHDNL  116
M_equina            RGNQKQGAQQKALAASIATFASVLVNKDAPLPDHLLSRIGHKHVSLGVTEDQYQIVHDNL  116
M_sympodialis       RGNQKQGAQQKALAASVATFASMLVNKDAPLPDHLLSRIGHKHVSLGVTEDQYQIVHDNL  116
M_caprae            RGNQKQGAQQKALAASVATFASMLVNKDAPLPDQLLSRIGHKHVSLGVTEDQYQIVHDNL  116
M_dermatis          RGNQKQGAQQKALAASVATFASMLVNKDAPLPDHLLSRIGHKHVSLGVTEDQYQIVHDNL  116
M_globosa           RGNQKQGAQQKALAASVATFASMLVNDDAPLPESMLSRIGHKHATVGVTEDQYQIVHDNL  111
M_restricta         RGNQKQGAQQKALAASVAVFASKLVSENEGLPAHLFSRIGHKHAALGITADQYQIVHDNL  113
                    *  *  *:* .*** ::   *  :           . ::.:**  : :  :.*  *   *

S_cerevisiae        LKAIKEVLG-DAATPEIINAWGEAYQAIADIFITVEKKMYEEAL---WPGWKPFDITAKE  158
M_yamatoensis       FGAIAKDLG-DAATPDIVEAWTEVYWLLARALINLERKLYAQQA--NNIVRAKFKLVKRT  162
R_nasimurium        FEAIAANLG-DILTPEIAEAWAEVYWLMADALIKLEKGLYASQA--NDVMFAPFRLVERK  162
K_kristinae         FGAIAKDLG-EAATEEVVAAWTEVYWLMADALIALEKGLYAVQA--NDVMRAPFRLVRRV  162
M_slooffiae         FGAIAKDLG-SAATPEVVEAWTEVYWLMANALIKLEKNLYAHQA--NNKIRAPFRVVSRK  162
B_ravenspurgense    FEAIVEVLGEDTVTPEVAEAWDAVYWIMARVLIDFEKDLYSTAGVEAGDVFRRVVVDKRE  180
M_vespertilionis    FGAIVHVLGADVVTADVAAAWTSVFWILANVLITFEKELYKGAGVQPGKVFRQAKVVDRD  171
M_cuniculi          FYAIVEVLGADVVTADVAAAWDSVYWIFARTLINFEKKLYAEVGAEPGKVFRTVSVVDRK  176
M_japonica          FGAIVEVLGADVVTQPVAEAWDDVFWIMARTLINMEKDLYSGAGVAPGKVFVSTRIVDRD  176
M_furfur            FGAIVAILGADVVTAPVAEAWENVYWIMANLLIKFEAELYEKAGVKPGEVFVNTQVVERK  176
M_obtusa            FGAIVAILGADVVTPPVAAAWEDVFWIMANLLIDFEAGLYTNAGVKPGDVFVNTRVLDRK  176
M_pachydermatis     FAAIVEVLGADVVTKDVAEAWDAVYWIMARVLIQFEKDLYKEAGVEPGKVWRSVKVAERR  176
M_nana_CBS9560      FYAIVQILGADVVTQDVAEAWDSVYWIMARVLIGFEKQLYEDAKVQPGQVFRQTTVVGRE  176
M_equina            FYAIVQILGADVVTQDVADAWESVYWIMARLLIDFEKDLYQGAKVEPGKVFCQTTVVDRV  176
M_sympodialis       FYAIVQVLGADVVTKDVAEAWDSVYWIMARLLINFEKDLYKGAKVEPGKVFRQTTVLERE  176
M_caprae            FYAIVQILGADVVTKDVAQAWNSVYWIMARLLINFEKDLYKGAKVEPGKVFRKTTVVERE  176
M_dermatis          FYAIVQILGADVVTKDVAEAWDSVYWIMARVLIDFEKDLYKGAKVEPGKVFRQTTVVERE  176
M_globosa           FAAIVQVLGADVVTKDVAEAWDRVFWIMGRLLIKFEKDLYAEAGVEPGKVFRQVKVVACD  171
M_restricta         FYAIVQVLGADTVTAEVAEAWDRVYWLMADMLIKFEKGLYNEAGVEPGKVFRQVKVVQSD  173
                    : **   ** .  *  :  **  .:  :.  :* .*  :*               :
```

*(continues in the next page)*

```
S_cerevisiae           YVASDIVEFTVKPKFGSGIELESLPITPGQYITVNTHPIRQENQYDALRHYSLCSASTK-  217
M_yamatoensis          QVTKDVVDMVFEPADNT-A---MTPGKAGQYISIYARTS---DGLLQPRQFTLLPSEET-  214
R_nasimurium           ETGENVIDLIFEPANAV-A---MTDAIAGQYVSIVTKAK---DGLRQARQFTLLPAEKN-  214
K_kristinae            RAADDVADLTFEPADDT-P---MTPAQAGQYVSIFCRAE---DGLLQPRQFTLLPSEPG-  214
M_slooffiae            ELGNHTADLTLERADDT-A---MTDAKPGQYVSAFAKAK---DGLLQPRQFTLLPCTKE-  214
B_ravenspurgense       DLSDTIVRFTVKAVDGS-A---LPAGRAGQYTSVGVALP---DGARQLRQYSLVASE---  230
M_vespertilionis       DRSANIVSFTVESTDAFNP---FPKHLAGQYVSVVSSLP---DGARQIRQYSLMDACEN-  224
M_cuniculi             DV-GDVAIFTVEGS----D---LPRHLPGQYISVGANLP---DGARQLRQYSLLDAGNA-  224
M_japonica             ERVGGIVRFSVESKDAAKP---LPAHQPGQYVSVRADLP---DGAHQLRQYSLIDSGVK-  229
M_furfur               DLSGGIVEFTVESKDASKP---LPAHQPGQYISVGAKLP---DGARQLRQYSLVDAGIN-  229
M_obtusa               ELGGGVVEFVVESKDASKP---LPVHQPGQYISVGAKLP---DGARQLRQYSLVDAGIK-  229
M_pachydermatis        GLVNGIVRFTIESTDASKP---LPTHQPGQYISVRAQLP---DGAGQLRQYSLLDAGVQ-  229
M_nana_CBS9560         GLTSDVVRFTIESKDSERP---LPAHLPGQYVSVRATLS---DGAGQLRQYSLIDAGNQ-  229
M_equina               NLTSDVVCFTIESKNGSQP---LPAHLPGQYISVRAKLP---DGAGQLRQYSLIDSGDK-  229
M_sympodialis          ALTSDVVRFTIESKDGSKP---LPAHLPGQYISVRAKLP---DGAGQLRQYSLVDSGEK-  229
M_caprae               SLTNDVARFTIESKDGSQP---LPAHLPGQYISVRAQLP---DGAGQLRQYSLIDSGKK-  229
M_dermatis             SLTSDVVRYTIESKDGSQP---LPAHLPGQYISVRAQLP---DGAGQLRQYSLIDSGEK-  229
M_globosa              TLAEGVTHFTIESTEASKP---LPGHKPGQYISVRAHLP---DGAGQLRQYSLTDAGKQS  225
M_restricta            KLTDDVTLFSIESTDASKP---LPAHKPGQYISVRARLP---DGAGQLRQYSLVDDGVK-  226
                                 .:                ***  :         :     *:::*

S_cerevisiae           -NGLRFAVKMEAARENFPAGLVSEYLHKDAKVGDEIKLSAPAGDFAINKELIHQNEVPLV  276
M_yamatoensis          --QRRI------AIKLDPHGEMTTIFQN-QEVGALLDISNPYGDMTLET-LETDPNSPLV  264
R_nasimurium           --QRRV------AIKLDTAGEMTPILHE-LQVGAIVEVSNPYGDVTLGG-FGDDESQPLY  264
K_kristinae            --QRRI------AVKLDPAGEMTPIILDKLTEGDVIEISNPYGDVTLGG-FGDDGDGPLW  265
M_slooffiae            --QRRI------AIKLDPAGEMTRIFLE-AHEGDVLELSNPYGDVTLGT-FGDDGKSPLY  264
B_ravenspurgense       EGKLSFVVKRVLAEADAPAGEVSNWLGDYAQVGTELDATVPFGDLVVD----IDGTAPVV  286
M_vespertilionis       PGQLRFAVKALQAQKDAPAGEVSNWLAKEVKVGTELAVSLPFGDLILD----TESQRPVV  280
M_cuniculi             GGRYVFAVKRVAAHDSAPEGEVSNWIWNNVKSGVEVEISLPFGELVLD----VQAESPVV  280
M_japonica             GSQLCFAVKALTETPDAPAGEVSNWLVQNAKVGDDLEISLPFGDLVLN----EEAQTPVV  285
M_furfur               PGRLTIAVKSVEATKDAPAGEVSNWLINNVKKGDDLEITLPFGDLVLN----EKSQRPVV  285
M_obtusa               PGQLTFAVKALEATGQAPAGEVSNWLIQNVKEGGDLEITLPFGDLVLD----QSSNCPVV  285
M_pachydermatis        AGRLSFAVKALEATHEQPAGEVSNWLIQNAQQGKELEVSLPFGDLVLD----TSSSSPVV  285
M_nana_CBS9560         PGQLSFAVKAVHGSETAPAGEVSNWLIENAQQGSNLEVSLPFGDFVLD----TQSESPVV  285
M_equina               PGHLRFAVKALHATESAPAGEVSNWLLENAQVGTDLEVSLPFGDLALN----TESSAPVV  285
M_sympodialis          PGRLSFAVKALEASESAPAGEVSNWLVDNAKVGTDLEVSVPFGDLVLN----TESNAPVV  285
M_caprae               AGGLSFAVKSLQASESAPAGEVSNWILENVQVGTDLEVSLPFGDLVLN----TESNAPVV  285
M_dermatis             AGRLSFAVKSLQATESAPAGEVSNWLLENAREGTDLEVSLPFGDLVLN----TESNAPVV  285
M_globosa              HGRLTFAVQAVKAHDQLPAGEVSNWLLENVKQGSELEISLPFGDLVLD----EQSNSPVV  281
M_restricta            AGRLSFAVKAVSATESAPAGEVSNWLSENARVGTELEVSLPFGDLVLD----TASSAPVV  282
                            .             * ::  : .    *  :  : * *:. :          *:

S_cerevisiae           LLSSGVGVTPLLAMLEEQVKCNPNRPIYWIQSSYDEKTQAFKKHVDELLAECANVDKIIV  336
M_yamatoensis          LICAGIGVTPVLAFVEKLAAQKSEREVMIIASSRSLAEAPLRGELLERAKELKKAKVLYG  324
R_nasimurium           LFSAGIGVTPMIAFLSELAATGSEREVVVVHADRSFTTWPLRDEMTAYVEMLENGRLVSF  324
K_kristinae            LFSAGIGVTPMIAFVSELVRQGSQREVTVVHTARDFASWPLREELAELVARLPHGRLISC  325
M_slooffiae            LIFAGIGVTPMLAFVHELAEQQSQRQVVVVGSAASKKEAPLYDELAAEVQRLPNGKLLFF  324
B_ravenspurgense       LLSAGIGTTPMMGILAALHAHASSREVVVVHCESSEEADVFAPERRALVDDLKGARLESI  346
M_vespertilionis       LISSGIGITPMFGMLARFVADRSSRKIVAIHCDQDMDGDAFYKERLHMVSQLSSGEAMTW  340
M_cuniculi             LISAGIGATPMIGMLSSLEAAKSQRTVLYLHAAASQEADTFASQRASLLEGIPNGKSEVW  340
M_japonica             LISAGIGVTPMLGMLSRLAETASSRRVISLHADKSASTDAFYEERKRLVSQLAQGSVDTW  345
M_furfur               LVSRGIGITPMLGILSYLAASNSSRPTLSLHADSSAATDAFFDERTKLVLKMPSGEAKTW  345
M_obtusa               LISAGIGVTPMLGILSHLAGTNSSRATLSLHADKSSATDVFFEERIQLISRLSSGEAKTW  345
M_pachydermatis        LISGGIGATPMIGMLSRLANDKSTRQVLVLHADHSEKTDALAAERHHLVSQLSNAKLNLF  345
M_nana_CBS9560         LISAGIGATSMIGMLSRLSVDASEHQVVVLHADTSSSTDALATERERLTSTIKNCEKHVF  345
M_equina               LISAGIGATPMMGMLSRLSSNASERKVIIVHADTSASTDAFATQREHLTSALKNCEKHIF  345
M_sympodialis          LISAGIGATPMIGMLSRLSSVASERHVVIVHADTNSGTDAFATHREHLASALKNYELHVF  345
M_caprae               LISAGIGATPMVGILSRLSTDASERRVVIIHADTSANTDAFAAQRDHLTAALKNCESHIF  345
M_dermatis             LVSAGIGATPMIGMLSRLSSDASERRVVIVHADTSASTDAFAAQRDHLTAALKNCESHIF  345
M_globosa              LISAGIGATPMISMLSRLALDKSEREVVVFHADSSAAADAFASQTKHLASQLPNCRFHTW  341
M_restricta            LVSAGIGATPMMGMLSRLVHDKSERQIIVLHVDKSAATDAFASQAKQLVSKLSNCRFHSW  342
                       *.  *:* * :..::         :    .    .      :  .
```

*(continues in the next page)*

```
S_cerevisiae          HTDTEPL------INAAFLKEKSPAHADVYTCGSLAFMQAMIGHLKELEHRDDMIHYEPF  390
M_yamatoensis         TTQEKDG--DFVG-RIDVSTLDIPANASVFLCGPLKFMQEMRSHLVEAGIAKHKIFYEIF  381
R_nasimurium          IEADGEG--DFEG-RVNVAELNVPANASAYLCGPLGFMQGVRSQLVEAGVPGTQIQYEIF  381
K_kristinae           TTASAEG--DHAG-RVRVPELDVPADAVAYLCGPLPFMKDVRSQLVDAGVPGQRIRYEIF  382
M_slooffiae           STKAQDG--EFSG-HVDVSKLQVPKEAIAYMCGPLPFMKDVRSQLVNAGVPGHQIQYEIF  381
B_ravenspurgense      YSDAA--------KRLSIDTLDLPRDAQVFVCGGTGFLEDARAQLRAAGFSDDAVNFELF  398
M_vespertilionis      HSVGEVTKDVKVG-RMDIASLTLPTDAQFYICGGTNFLKSICDGLRKIHVPEEDLHFELF  399
M_cuniculi            YSHGPGAR---SG-HIDLSQVELPGSAEFYLCGGDAFLKDMRTALDENGIPSSKVHFELF  396
M_japonica            YTDGDAQTEVNQG-RFDLSSQSVPADAEYYLCGANEFLQSLRTTLNAWDVPKDRVHFELF  404
M_furfur              YSEGQTGPNVSQG-HLDLSTVQLPADAEIYLCGGNEFLQRMRKQLKDMKVPDERVYFELF  404
M_obtusa              HAEDESGSDVLKG-RLDLSAIQLPSEAQYYLCGGNEFLQSMRKQLQELNVPADRVNFELF  404
M_pachydermatis       YSHGEAGPHITVS-HLDLKHVELPHDAQFYLCGSTDFLQAARESLKSMNIDAESVHFELF  404
M_nana_CBS9560        YHEGAANENTTIA-RMDLSKLEFPEGAEFYLCGSTSFLQEAREGLKANRVEPTSVHYELF  404
M_equina              YKEGTASNGSTIG-QLQVGQLTFPEHAEFYLCGSLTFLQEAREGLKAKNVSPSAVHFELF  404
M_sympodialis         YKEGASGKNVTIG-HLDVSKFAFPNDAEFYLCGSTTFLQEAREGLKANNVKPSSVHFELF  404
M_caprae              YKEGAPGKDVTIG-QLDVSKFSFPNDAELYLCGSTTFLQEAREGLKVNNVKSSSVHFEMF  404
M_dermatis            YKEGASGKDVTVG-HLDVSKLTFPNDAEFYLCGSTAFLQEAREGLKANNVKPSSVHFELF  404
M_globosa             YAQGESNETTTVGDRLDLTKETLPSNAQYYLCGSTAFLQACSKDLESLSVPASHTHYELF  401
M_restricta           YAEGTSDDTTTIGEKLNLGKESLPKDALVYLCGSTEFLQAAAKDLKETGIPEDNVQFELF  402
                                    .  .     *  *  : **   *::     *           :* *

S_cerevisiae          GPKMSTVQV---------      399
M_yamatoensis         GTDQWMLHDQNRSE----      395
R_nasimurium          GPDQWMLHDQARA-----      394
K_kristinae           GPDQWMLHDQAREEPAAA      400
M_slooffiae           GPDQWMLHDQARE-----      394
B_ravenspurgense      APNDWLLDG---------      407
M_vespertilionis      TPNDWLL-----------      406
M_cuniculi            SPNDWLIDY---------      405
M_japonica            TPNDWLLDA---------      413
M_furfur              SPNDWLLDE---------      413
M_obtusa              SPNDWLLDV---------      413
M_pachydermatis       APNDWLLDE---------      413
M_nana_CBS9560        MPNDWLLDE---------      413
M_equina              SPNDWLLDE---------      413
M_sympodialis         MPNDWLLDE---------      413
M_caprae              APNDWLLDE---------      413
M_dermatis            TPNDWLLDE---------      413
M_globosa             TPNEWLLK----------      409
M_restricta           TPNDWLIS----------      410
                        .   :
```

**Fig. S12.** Structural features depicted on the amino acid sequence alignment of the Yhb1 and Yhb101 flavohemoglobins. Residues shaded in brown represent the linker region separating the N-terminal globin and C-terminal oxidoreductase domains. Amino acids that surround the active site or are known to play a role in catalysis are highlighted in blue; these include Y29 responsible for $O_2$ binding, and other uncharged residues that surround the heme and provide for a relatively hydrophobic active site (Q53, L57, V61 or I61). Highlighted in purple are the amino acids H85, Y95, Y126, E137, and Y141, which are known to form an H-bonding network; note in dark purple the non-conserved residue F126. Within the oxidoreductase domain, side chains responsible for FAD binding are shown in green (G187-Y189, and R206-S209), and those responsible for NADPH binding are in yellow (G280-T285, and A357-G373); in dark green and dark yellow are highlighted different residues specific of the flavohemoglobin Yhb101. Amino acid numbering is based on the *S. cerevisiae* Yhb1 sequence. In the bottom line, asterisks indicate conserved amino acid residues, colon (:) indicates conservation between groups of strongly similar properties (scoring > 0.5 in the Gonnet PAM 250 matrix), and period (.) indicates conservation between groups of weakly similar properties (scoring =< 0.5 in the Gonnet PAM 250 matrix). Most of the known functional regions are conserved in both the

*Malassezia* Yhb1 and Yhb101, with the notable differences being the lack of conserved tyrosine (Y126) residues in the hydrogen bond in *M. vespertilionis, M. japonica, M. obtusa* and *M. globosa.* Other variations are also found in the cofactor binding sites of the reductase domain, in particular, 1) the isoalloxazine FAD binding site was different between the two *Malassezia* flavohemoglobin, with RQYS being the Yhb1 motif, and RQFT being that of the Yhb101, which is also shared with the putative donor bacteria (*K. kristinae*); and 2) the nicotinamide adenine dinucleotide (NADH) motif being GIGXT for Yhb1, and GIGVT for Yhb101.

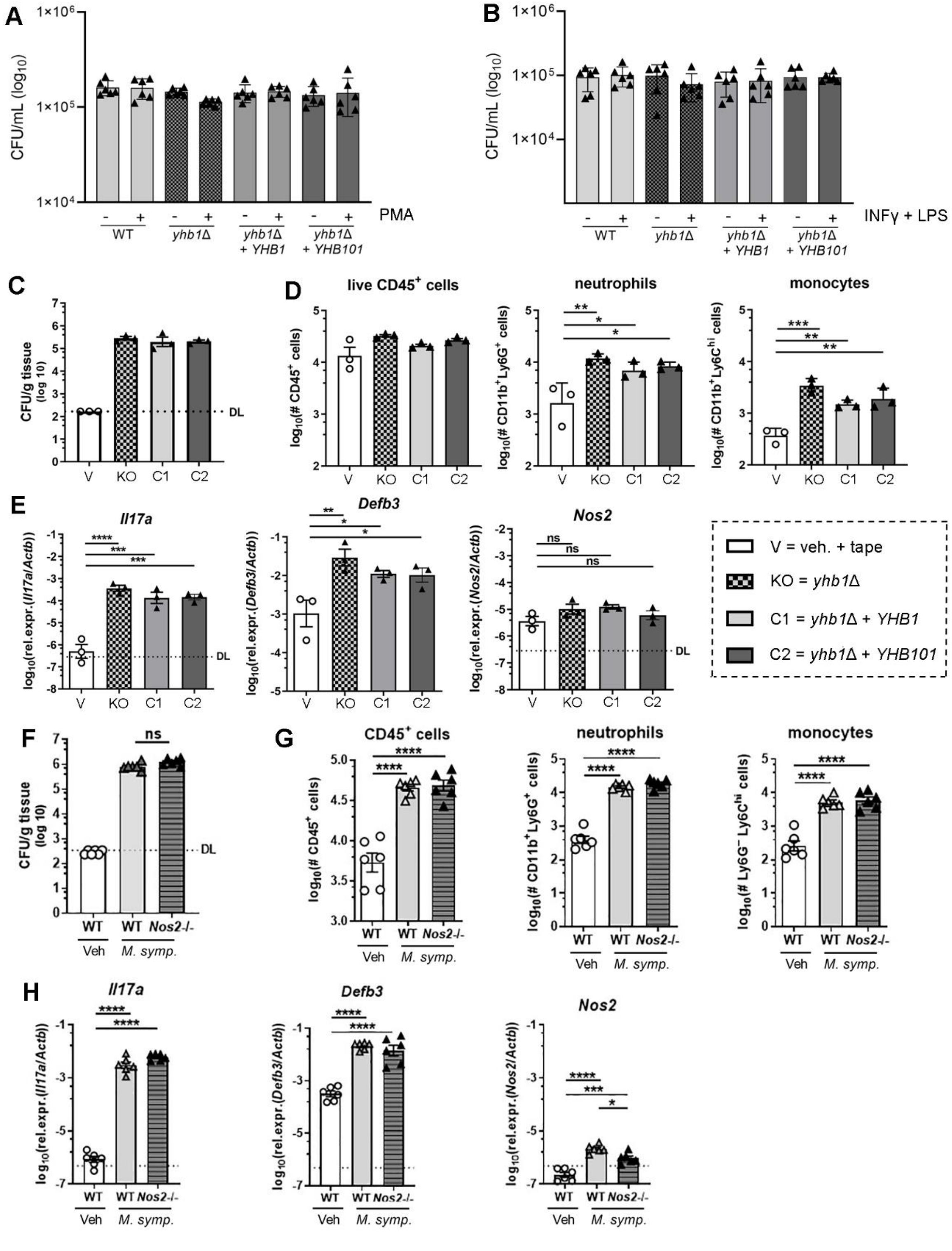

**Fig. S13.** ***Malassezia*** **flavohemoglobins are not required for pathogenesis in** ***ex vivo*** **and** ***in vivo*** **models.** (A-B) *M. sympodialis* WT, *yhb1*Δ, *yhb1*Δ + *YHB1*, and *yhb1*Δ + *YHB101* were opsonized with 20% human serum and added to J774A.1 macrophage (1:1 ratio) activated with phorbol myristate acetate (PMA) (A) or interferon-gamma +

lipopolysaccharide (IFNγ + LPS) (B). After 24 h, J774A.1 cells were lysed and *M. sympodialis* cells were harvested and plated on selective media to determine CFUs. Data are expressed as log10 of colony forming units (CFU)/mL. (C-E) WT C57BL/6j mice were infected epicutaneously with *M. sympodialis* WT, *yhb1*Δ, *yhb1*Δ + *YHB1*, and *yhb1*Δ + *YHB101*. Skin colony forming units (CFU) (C), cellular infiltrates (D) and *Il17a*, *Defb3* and *Nos2* transcripts (E) were analyzed on day 4 post infection. (F-H) WT and *Nos2*-/- mice were infected epicutaneously with *M. sympodialis* WT. Skin CFU (F), cellular infiltrates (G) and *Il17a*, *Defb3* and *Nos2* transcripts (H) were analyzed on day 5 post infection. Graphs show the mean + SEM of each group. Individual samples are also indicated with each symbol representing one well (A-B) or one mouse (C-H). The dotted line indicates the detection limit. Data are from one experiment (C-E) or pooled from two independent experiments (A-B, F-H). Statistics are calculated using one-way ANOVA with Sidak's multiple comparisons test. *$p<0.05$, **$p<0.01$, ***$p<0.001$, ****$p<0.0001$.

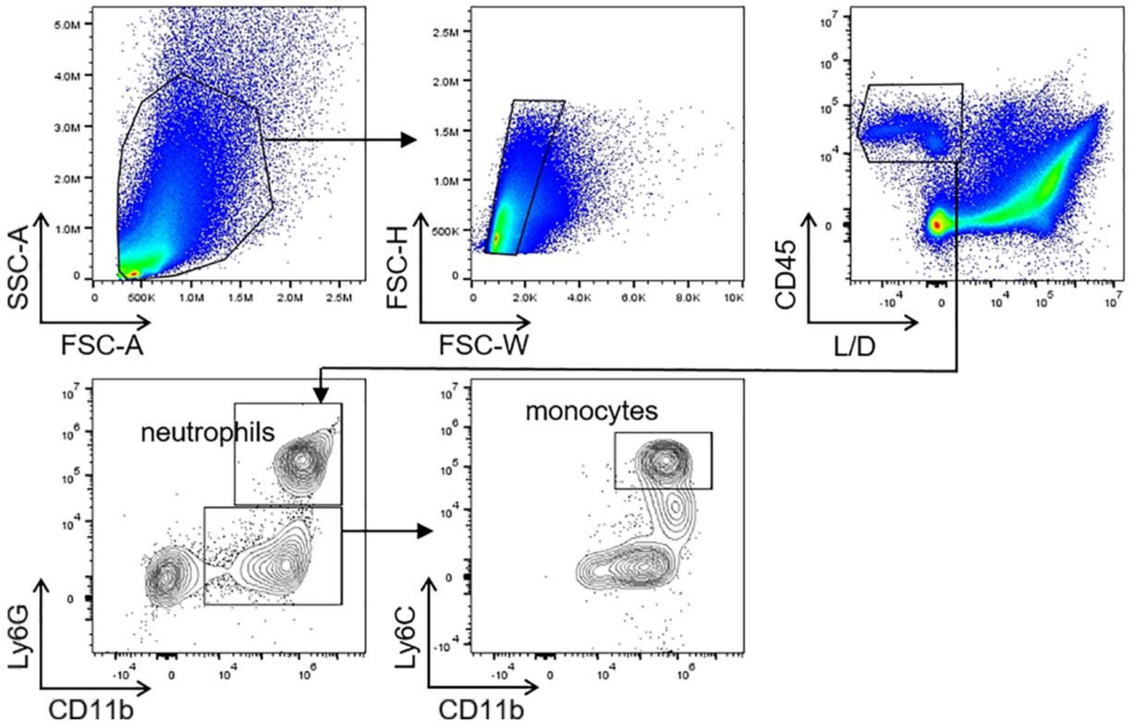

**Fig. S14:** Gating strategy for myeloid cells in the ear skin. Cells were pre-gated as scatter$^+$, singlets, viable and CD45$^+$. Within the CD45+ population, CD11b$^+$ Ly6G$^+$ cells were identified as neutrophils and CD11b$^+$ Ly6G$^-$ Ly6C$^{hi}$ cells as monocytes.

**Supplementary tables**

**Table S1.** Scaffold/chromosomal locations, and accession numbers of genomic regions surrounding the *Malassezia* flavohemoglobin-encoding genes used for synteny comparisons displayed in Figure 2B.

| Species | Strain | Scaffold/Chromosome | Coordinates | GenBank Accession |
|---|---|---|---|---|
| *Malassezia caprae* | CBS10434 | LFFV01000091.1 | 147,064..115,068 | GCA_001264625.1 |
| *Malassezia dermatis* | CBS9169 | LFFX01000100.1 | 247,963..279,479 | GCA_001264665.1 |
| *Malassezia sympodialis* | ATCC42132 | LT671826.1 | 332,514..300,264 | GCA_900149145.1 |
| *Malassezia equina* | CBS9969 | LFFY01000032.1 | 211,400..243,389 | GCA_001264685.1 |
| *Malassezia nana* | CBS9557 | Scaffold_4 | 297,384..328,773 | GCA_001265015.1 |
| *Malassezia pachydermatis* | CBS1879 | NW_017253143.1 | 316,054..349,953 | GCF_001278385.1 |
| *Malassezia globosa* | CBS7966 | NW_001849865.1 | 454,282..489,698 | GCA_000181695.1 |
| *Malassezia restricta* | KCTC27527 | scaffold00006 | 398,082..429,794 | GCA_003290485.1 |
| *Malassezia furfur* | CBS14141 | Chr2 | 1,384,981..1,422,441 | GCA_009938135.1 |
| *Malassezia yamatoensis* (*YHB101* region) | MY9725 | LFCX01000037.1 | 15,445..1 | GCA_001264885.1 |
| *Malassezia yamatoensis* | MY9725 | LFCX01000042.1 | 1,006,753..977,400 | GCA_001264885.1 |
| *Malassezia obtusa* | CBS7876 | LFGC01001518.1 | 21,446..41,670 (end) | GCA_001264985.1 |
| *Malassezia japonica* | JCM11963 | Scaffold_1 | 958,498..927,858 | GCA_001600795.1 |
| *Malassezia vespertilionis* | CBS15041 | KZ454991.1 | 571223..603556 | GCA_002818225.1 |
| *Malassezia cuniculi* | CBS11721 | LFFW01000003.1 | 302,043..335,709 | GCA_001264635.1 |
| *Malassezia slooffiae* (*YHB101* region) | CBS7956 | YHSBHY01000006.1 | 83,812..109,097 | GCA_010577765.1 |
| *Malassezia slooffiae* | CBS7956 | YHSBHY01000005.1 | 379,150..345,760 | GCA_010577765.1 |

**Table S2.** Primers used in the present study.

| **Name** | **Sequence (5' - 3')** | **Purpose** | **Note** |
|---|---|---|---|
| JOHE44264 | GCGCGCCTAGGCCTCTGCAGGTCGACTCTGTG TGAACCAATAGAGTCC | *M. sympodialis yhb1*Δ - F 5' flanking region | Underlined is the chimeric region for recombination in plasmid pGI3 |
| JOHE44265 | **GAGGATCTGCACCGTGGA**ATCACTTCACCGT GTTCTGG | *M. sympodialis yhb1*Δ - R 5' flanking region | In bold is the region for recombination with JOHE43277/ALID2078 |
| JOHE44266 | **CAGACATAGGAGAGGACG**TTCAGGAAGCTC GCGAAGG | *M. sympodialis yhb1*Δ - F 3' flanking region | In bold is the region for recombination with JOHE43279/ALID2081 |
| JOHE44267 | TGATTACGAATTCTTAATTAAGATATCGAGAA TCCTCAGACAATTCGTCG | *M. sympodialis yhb1*Δ - R 3' flanking region | Underlined is the chimeric region for recombination in plasmid pGI3 |
| JOHE43277 | TCCACGGTGCAGATCCTC | *Malassezia NAT* - F | |
| JOHE43278 | CGTCCTCTCCTATGTCTG | *Malassezia NAT* - R | |
| JOHE44268 | TCAATCAATGAACATATAGG | *M. sympodialis yhb1*Δ identification - F | |
| JOHE44269 | GAGGAATCTGAAGATGAGC | *M. sympodialis yhb1*Δ identification - R | |
| JOHE43281 | GTCGGAGAAGCAGTCAATGC | *M. sympodialis yhb1*Δ identification - R | Designed on the gene marker |
| JOHE43282 | CACCAGGGTTTCCAGTCTC | *M. sympodialis yhb1*Δ identification - F | Designed on the gene marker |
| JOHE43279 | CTGATCCAAGCTCAAGCTC | Assess correct recombination in pGI3 - F | |
| JOHE43280 | GTTGGCCGATTCATTAATGC | Assess correct recombination in pGI3 - R | |

| | | | |
|---|---|---|---|
| JOHE45236 | GCGCGCCTAGGCCTCTGCAGGTCGACTCTGAT GTTGAAGAGACGACCAG | *M. sympodialis* promoter and *YHB1* gene - F | Underlined is the chimeric region for recombination in plasmid pGI3 |
| JOHE45237 | GCCTCCGCCTCCGCCTCCTTCGTCAAGAAGCC AGTCGT | *M. sympodialis* promoter and *YHB1* gene - R | In red is the glycine linker |
| JOHE44555 | GGAGGCGGAGGCGGAGGCATGGTGAGCAAG GGCGAGGA | GFP - F | In red is the glycine linker; in green the region for GFP |
| JOHE45235 | ACACATCACTTGGATTCCACTAGTACAGCTCG TCCATGCC | GFP - R | In green the region for GFP; underlined is the region for recombination with the terminator |
| JOHE44557 | GGCATGGACGAGCTGTACTAGTGGAATCCAA GTGATGTGT | terminator - F | Underlined is the region for recombination with GFP |
| JOHE44549 | **GAGGATCTGCACCGTGGA**AAGTCGTCGGGG CTGAATC | terminator - R | In bold is the region for recombination with JOHE44550 |
| JOHE44550 | TCCACGGTGCAGATCCTC | NEO - F | |
| JOHE44551 | TGATTACGAATTCTTAATTAAGATATCGAGCG TCCTCTCCTATGTCTG | NEO - R | Underlined is the chimeric region for recombination in plasmid pGI3 |
| JOHE99961 | GCGCGCCTAGGCCTCTGCAGGTCGACTCTTGA TAGTGTCTATTATTAGG | *M. yamatoensis* promoter and *YHB101* gene - F | Underlined is the chimeric region for recombination in plasmid pGI3 |
| JOHE99962 | GCCTCCGCCTCCGCCTCCGGACCGATTCTGAT CGTGG | *M. yamatoensis* promoter and *YHB101* gene - R | In red is the glycine linker |
| JOHE45289 | TTCACGCACATACCTGCAAT | *M.nana YHB1* amplification - F_1 | F_1 was combined both with R_1 and R_2 |
| JOHE45290 | GCGAGCAACAAGAAAAGGAC | *M.nana YHB1* amplification - F_2 | F_2 was combined both with R_1 and R_2 |

| | | |
|---|---|---|
| JOHE45291 | TGACGCATCCACTGAGAGAC | *M.nana YHB1* amplification - R_1 |
| JOHE45292 | TCAAGAAAATGCTGCACAGG | *M.nana YHB1* amplification - R_2 |
| JOHE45545 | CAGACAACACAGCAATGACACC | *M. yamatoensis YHB101* F - qPCR |
| JOHE45546 | CGTTGAGTCTCCTCAGATGG | *M. yamatoensis YHB101* R - qPCR |
| JOHE45547 | GTCATTGGGTGTGACAGAG | *M. sympodialis YHB1* F - qPCR |
| JOHE45548 | AGACGCGCCATAATCCAGTAT | *M. sympodialis YHB1* R - qPCR |
| JOHE45549 | TGCCGGAGCTCACCTCGCA | *M. sympodialis - M. yamatoensis TUB2* F - qPCR |
| JOHE45550 | TACGACGAGTTCTTGGTCTG | *M. sympodialis - M. yamatoensis TUB2* R - qPCR |
| CID101552_PCR1_For | TGTTTAACTTTAAGAAGGAGATATACCATGGGCCATCACCACCATCACCAC | N-term 8XHis-Tev tag |
| CID101552_PCR1_Rev | ATAACTGGTTGGGACTTGGTGCTTAGACCGCTACCCTGGAAATACAGATTTTC | N-term 8XHis-Tev tag |
| CID101552_PCR2_For | CTAAGCACCAAGTCCCAACCAGTTAT | *M. yamatoensis YHB101*coding region for protein expression |
| CID101552_PCR2_Rev | GTGGTGGTGCTCGAGTGCGGCCGCAAGCTAAGCTTATCATTCGGACCGATTCTGATCGTGGAGC | *M. yamatoensis YHB101* coding region for protein expression |
| IL17A_1 | GCTCCAGAAGGCCCTCAGA | murine *Il17A* - F |
| IL17A_2 | AGCTTTCCCTCCGCATTGA | murine *Il17A* - R |

| | | |
|---|---|---|
| Defb3_1 | GTCTCCACCTGCAGCTTTTAG | murine *Defb3* - F |
| Defb3_2 | ACTGCCAATCTGACGAGTGTT | murine *Defb3* - R |
| Nos2_1 | GTTCTCAGCCCAACAATACAAGA | murine *Nos2* - F |
| Nos2_2 | GTGGACGGGTCGATGTCAC | murine *Nos2* - R |
| Actb_1 | CCCTGAAGTACCCCATTGAAC | murine *Actb* - F |
| Actb_2 | CTTTTCACGGTTGGCCTTAG | murine *Actb* - R |

**Table S3.** Data collection, phasing and refinement statistics relative to the X-ray structure of *M. yamatoensis* flavohemoglobin Yhb101

| | Anomalous (Fe phasing) | Native (refinement) |
|---|---|---|
| **Data collection** | | |
| Space group | $P2_12_12$ | $P2_12_12$ |
| Cell dimensions | | |
| *a, b, c* (Å) | 91.08, 110.68, 39.01 | 91.08, 110.68, 39.01 |
| $\alpha, \beta, \gamma$ (°) | 90, 90, 90 | 90, 90, 90 |
| Resolution (Å) | 50.0 – 1.70 (1.74-1.70) | 50.0 – 1.70 (1.74-1.70) |
| $R_{sym}$ or $R_{merge}$ | 5.7 (54.6) | 5.7 (54.2) |
| $I / \sigma I$ | 24.66 (2.46) | 29.07 (3.12) |
| Completeness (%) | 99.9 (98.8) | 99.9 (98.5) |
| Redundancy | 9.5 (4.4) | 13.5 (6.4) |
| **Refinement** | | |
| Resolution (Å) | | 47.26-1.70 (1.74-1.70) |
| No. reflections | | 44,179 (4297) |
| $R_{work}$ / $R_{free}$ | | 0.167/0.198 (0.240/0.271) |
| No. atoms | | |
| Protein | | 2998 |
| Ligand/ion | | 104 |
| Water | | 428 |
| *B*-factors | | |
| Protein | | 24.29 |
| Ligand/ion | | 18.30 |
| Water | | 34.04 |
| R.m.s deviations | | |
| Bond lengths (Å) | | 0.006 |
| Bond angles (°) | | 0.91 |

*Data was collected on a single crystal.
*Values in parentheses are for highest-resolution shell.

**Additional Datasets**

**Dataset S1 (separate file).** Upregulated differentially expressed genes for FDR <0.05 in the condition *M. sympodialis yhb1*Δ versus *M. sympodialis* WT. Highlighted in yellow are DEGs with $log_2$FC < 0.5.

**Dataset S2 (separate file).** Downregulated differentially expressed genes for FDR <0.05 in the condition *M. sympodialis yhb1*Δ versus *M. sympodialis* WT. Highlighted in yellow are DEGs with $log_2$FC < - 0.5.

**Dataset S3 (separate file).** Upregulated differentially expressed genes for FDR <0.05 and $log_2$FC ≥ 0.5 in the condition *M. sympodialis* WT + $NaNO_2$ versus *M. sympodialis* WT untreated.

**Dataset S4 (separate file).** Downregulated differentially expressed genes for FDR <0.05 and $log_2$FC ≤ -0.5 in the condition *M. sympodialis* WT + $NaNO_2$ versus *M. sympodialis* WT untreated.

**Dataset S5 (separate file).** Summary of the HGT candidate genes identified in *Malassezia* species. Each sheet is indicated with a color that matches the *Malassezia* phylogeny represented in Figures 2A and 6.
